# A positive feedback loop of ‘MPK3-PIN1A trafficking-auxin flux’ trio governs dual gravitropic and wounding response in rice

**DOI:** 10.1101/2025.02.12.637814

**Authors:** Mrinalini Manna, Balakrishnan Rengasamy, Sarvesh Jonwal, Uttam Pal, Gopal Banerjee, Alok Krishna Sinha

## Abstract

The auxin flow in plants plays a pivotal role in gravitropic organ movement and tissue regeneration following injury. However, the regulatory aspects of these phenomena are not entirely understood. In this study, we found that hyperactivity of MAP Kinase 3 (MPK3) led to downregulation of *PINs* and reduction in tissue auxin content, impairing gravitropic response of rice and accelerating tissue senescence following mechanical wounding. The MPK3 was found to phosphorylate PIN1A majorly at its Ser^351^ residue, mutation of which into its phospho-mimic variant enhanced latter’s endosomal trafficking and consequently improved auxin flux. Further, overexpression of the phospho-null variant of *PIN1A* lowered gravitropic response of rice, thus causing wider tiller angle similar to the *pin1a* knockout lines. Additionally, upon wounding, their tissue senescence was faster in comparison to the overexpression lines of wild-type and phospho-mimic variants of *PIN1A*, which also displayed improved resistance to wounding by *Bipolaris oryzae*. The PIN1A trafficking-mediated auxin flux also regulated MPK3 activity via a positive feedback loop. Together, these results uncover a novel feedback loop encompassing “MPK3-PIN1A trafficking-auxin flux” trio that regulates dual gravitropic and wounding response in rice.

## Introduction

Auxin flux throughout the plant system sustains its life and is a vital contributor of tissue differentiation (Su et al., 2011), organogenesis (Suzuki et al., 2023), plant movement (Tivendale and Millar, 2022), tissue regeneration (Canher et al., 2020; Hoermayer et al., 2020), etc. Among various auxin transporters present in plants, PIN auxin efflux transporters constitute the vital component of apoplastic directional flow of auxin (Manna et al., 2022; Ung et al., 2022) In recent years, various protein kinases including MAP kinases have been implicated in the control of PIN activity, polarity and trafficking (Lanassa Bassukas et al., 2022). Phosphorylation of the hydrophilic loop domain of PINs by PINOID and MAPKs are known to aid protein localization in Arabidopsis. However, even after a century of passionate exploration, the mysteries of auxin biosynthesis, transportation, metabolism and signaling have not been completely understood (Yu et al., 2022). The present study ensigns upon how PIN1A trafficking regulates differential auxin distribution to control shoot gravitropism or tiller angle in rice and maintain its tissue integrity following an injury.

In rice, ideal tiller angle is of economic significance as neither spreading nor compact plant architecture is useful for obtaining maximum yield. Very compact plant architecture favors disease development while spreading architecture reduces grain yield per unit area as plant density gets reduced (Wang et al., 2022). Earlier studies have shown that shoot gravitropism is strongly associated with tiller angle in rice (Huang et al., 2021; Li et al., 2021) and dampening of shoot gravitropism leads to wider tiller angles (Huang et al., 2021; Xia et al., 2021). Despite momentous advancement in understanding the molecular basis of shoot gravitropism in Arabidopsis, the regulatory network that maintains shoot gravitropism and thus tiller angle in rice is not well understood. We discovered an important auxin flux-mediated regulatory switch that maintains tiller angle in rice. Apart from maintaining shoot gravitropism, auxin flux is known to play a central role in maintaining tissue regenerative processes, such as recovery from wounding, tissue damage or organ loss (Melnyk, 2016; Xu, 2018). Canher et al. (2020) showed how vascular stem cells obstruct polar auxin flux, much similar to rocks in a stream thus causing auxin accumulation around the periphery of damaged tissue which in turn facilitates periclinal cell division in endodermis causing replenishment of vascular stem cell pool. In the present study, we have demonstrated how PIN1A trafficking controls the spread of necrotic lesions following tissue injury.

Many cellular and physiological processes operate in a circular, self-regulating manner to maintain homeostasis. The self-operating nature of life is essentially determined by feedback mechanisms which can either be a positive feedback loop or a negative feedback loop. Positive feedback loops amplify the original stimulus. For example, upon sensing lower ambient nutrient levels, plants uptake more water which in turn speeds up hydraulic flux-mediated auxin circulation (Mehra et al., 2022) resulting in more root branching and faster plant growth, thus setting up a condition to uptake more water along with dissolved nutrients to sustain the speedy growth of plants (Manna et al., 2024a). In contrast, negative feedback loops counteract changes to bring a system back to a set point. For example, when cellular auxin concentrations are low, Aux/IAA proteins act as repressors to inhibit DNA-binding ARF (Auxin Response Factor) transcription factors from binding to the auxin response elements (AuxREs) present in the promoters of auxin-inducible genes including *IAAs* thus repressing their transcription (Qi et al., 2024). There are many such feedback loops in auxin signaling pathway still unexplored. The present study has discovered one such positive feedback loop that regulates dual tiller angle and injury mediated tissue senescence in rice.

## Results

### The nexus of auxin biosynthesis, its distribution and MAP kinase activity determine organ curvature in rice

Roots don’t grow straight, as there are bents and turns along their entire length (Kircher and Schopfer, 2016). Observing one such bent region in the root of a *DR5-gus* expressing rice seedling revealed more gus staining in the convex side of curvature (Figure 1A). This region undoubtedly accumulated more auxin that favored the emergence of lateral roots invariably from the convex side of the curved portions of the primary roots (Figure 1A, B, E). Further, differential auxin distribution caused greater cell expansion in the convex side causing roots to form the curvature (Figure 1C), while a strait growing root had similar cellular expansion (Figure 1D). The bending of a plant organ falls under the category of gravitropic response (Luschnig and Friml, 2024). To understand the role of auxin in regulating the gravitropic response of rice seedlings, we analyzed the extent of shoot bending in the absence or presence of external auxin (here NAA; 1-naphthaleneacetic acid). The study revealed no significant difference in extent of shoot curvature (Figure 1F, G) when rice roots had usual or even higher auxin content (Figure 1H). An increase in the auxin level of rice roots occurred due to combined effect of increased auxin uptake from the medium as well as upregulation of auxin biosynthesis pathway genes, *TARs* and *YUCCAs* (Manna et al., 2022; Figure 1P). Consequently, the rice roots were found to have lesser MAP Kinase (MPK; here, MPK3 and 6) activity when auxin content of the tissue was higher (Figure 1I) along with having lesser transcript abundance of *MPK3* (Figure 1J; since further study encompasses MPK3 signalling, we shall focus upon MPK3 only hereafter). Interestingly, in the presence of TIBA (Tri-Iodobenzoic Acid; an auxin transport inhibitor), gravitropic response of the rice seedlings got dampened as the shoots couldn’t bend as quickly as the ones grown without TIBA (Figure 1K, L). These seedlings had lower auxin content (Figure 1M) and higher MPK3 activity in roots despite having a visible reduction in transcript abundance of *MPK3* (Figure 1N, O). TIBA treatment also downregulated the expression of auxin biosynthesis pathway genes in the root causing reduction in auxin level (Figure 1P). Additionally, we observed that both NAA and TIBA treatment resulted in downregulation of *PIN* class of auxin transporters (Figure 1Q). When auxin was externally supplemented, PIN mediated lower auxin transportation thus couldn’t affect gravitropic response of rice seedlings as abundant supply of auxin could nullify the effect of decreased auxin transportation required for cell expansion and organ bending. On the other hand, when both auxin biosynthesis and its transportation were disrupted in the presence of an auxin transporter inhibitor, shoot bending was deeply affected. This revealed the importance of PIN-mediated auxin transportation in the plant’s gravitropic response. Furthermore, the study also pointed out that MPK3 activity might have a bearing on the gravitropic response of the rice seedlings.

**Figure 1.**
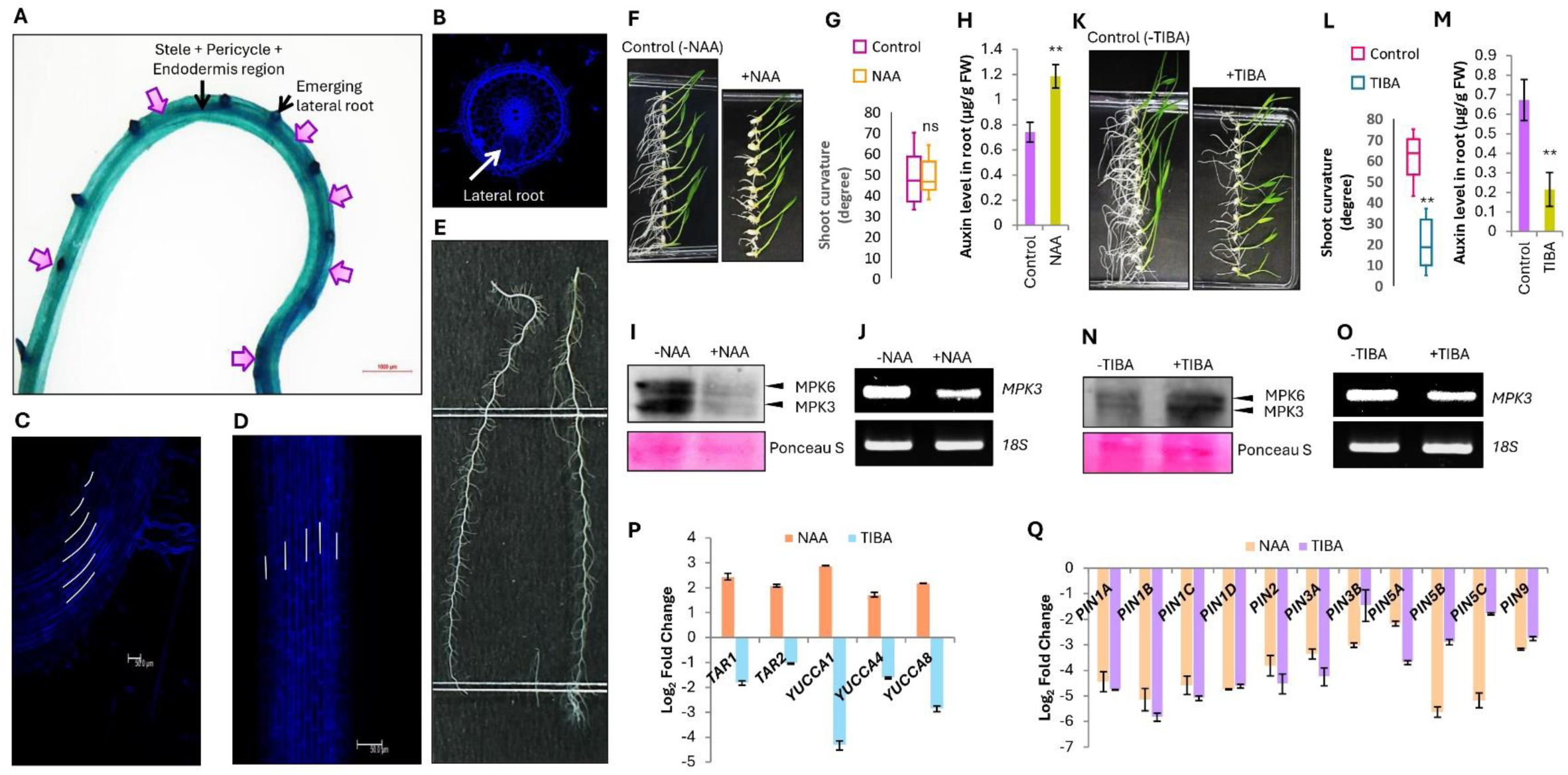
Resolving the cellular activities when a plant organ bends. **(A)** *DR5-gus* expressing curved portion of a root showing more auxin accumulation in the convex side which results in lateral root formation **(B)** from this region. The curvature is due to more cell expansion in the convex side **(C)** as similar cell expansion results in straight growth of root **(D)**. **(E)** The entire primary roots of a rice seedling showing presence of curvatures and lateral roots invariably arising from the convex sides highlighting the significance of auxin built-up in causing curvature. External auxin (NAA) supplementation did not dampen gravitropic response of rice seedlings (n = 10) **(F, G)** when there was increase in root auxin level (n = 8) **(H)**, lower MPK3 activity **(I)** and downregulation of *MPK3* expression **(J)**. But, TIBA supplementation led to dampening of their gravitropic response (n = 10) (**K, L**), lower root auxin level (n = 8) **(M)**, increased MPK3 activity **(N)** despite a downregulation in its expression **(O)**. **(P)** Further, auxin supplementation led to upregulation of auxin biosynthesis genes; *TARs* and *YUCCAs* while TIBA treatment led to their downregulation (n = 3). **(Q)** Both NAA and TIBA treatment caused downregulation of *PIN* genes’ expression (n = 3).

### MPK3 activity regulates PIN and auxin abundance to control gravitropic response

To understand the role of MPK3 activity on rice seedling’s capacity to bend, we investigated the gravitropic response of the DEX inducible *MPK3* overexpression lines (Jonwal et al., 2023; Singh et al., 2019) as well as an *MPK3* knockout line (Supplementary Figure S1A-S1F). While mock (here DMSO) treatment had no significant difference in the extent of shoot curvature, DEX treatment dampened the gravitropic response of the *MPK3* overexpression lines (Figure 2A-D). Interestingly, we found that overexpression of *MPK3* slowed rice seedling’s growth (Supplementary Figure S2A-S2E), lowered auxin level (Supplementary Figure S2F) and downregulated *PIN* expression in the rice roots (Supplementary Figure S2G). On the other hand, knockout of *MPK3* had opposite effects on rice seedlings as evidenced by greater root elongation (Supplementary Figure S3A-S3D), increased auxin content (Supplementary Figure S3E) and upregulation of *PIN* genes in the root tissue (Supplementary Figure S3F). It was further discovered that external supplementation of NAA could restore gravitropic response of the *MPK3* overexpression lines (Figure 2E-G). The study hence proved that an increase in MPK3 activity affects *PIN* abundance, which in turn reduces auxin flux (the same as happens in the case of TIBA treatment) in rice that ultimately metamorphoses into the dampened gravitropic response. Further, auxin supplementation overcomes this effect by restoring auxin supply to the regions undergoing cell expansion for bending.

**Figure 2.**
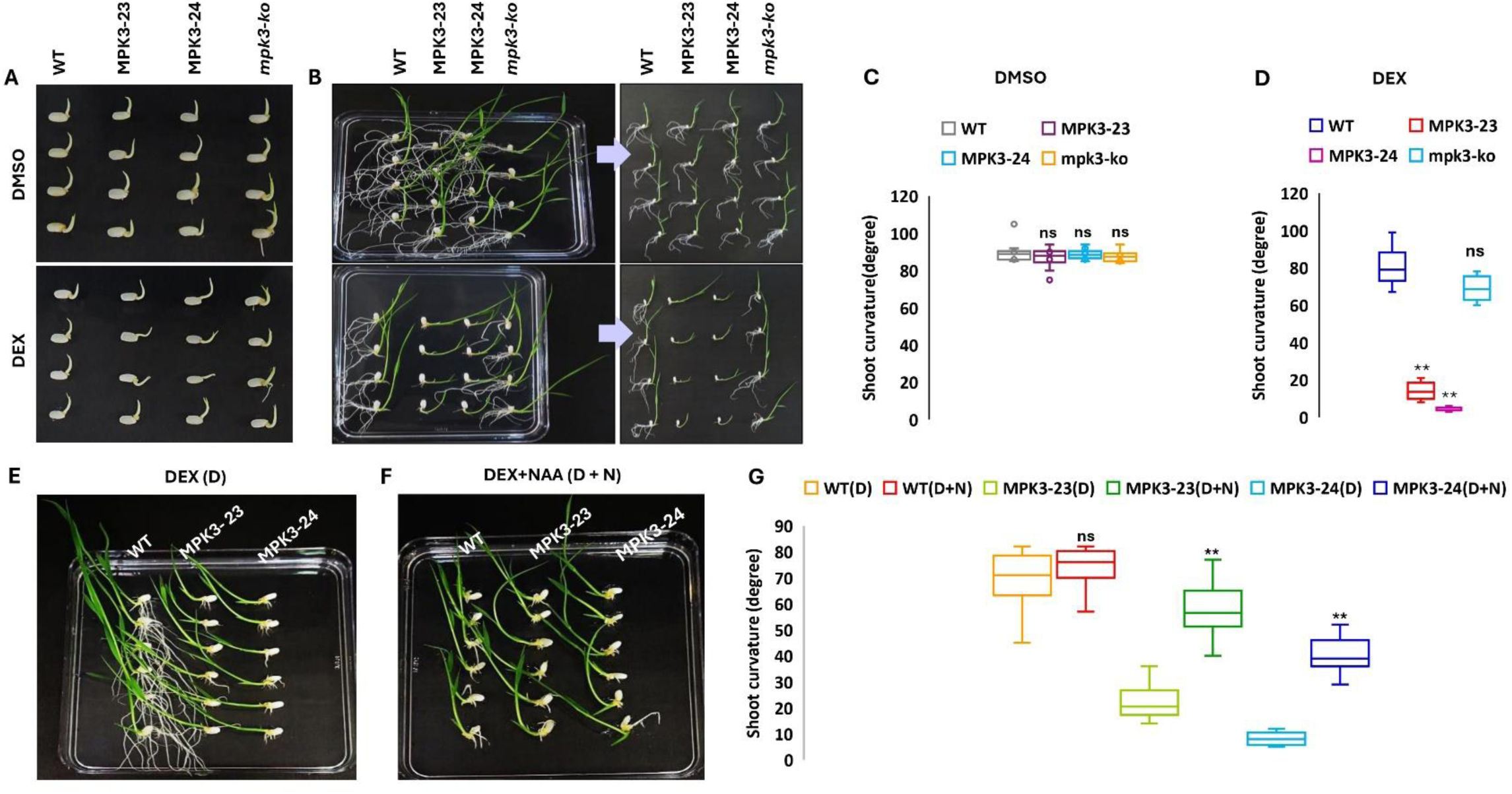
Comparison of gravitropic response of WT plants, DEX inducible *MPK3* overexpression lines and an *mpk3* knockout line. While there was uniform gravitropic response at seed germination **(A)** and seedling **(B, C)** stages of various rice plants during mock (DMSO) treatment (n = 10), DEX treatment dampened gravitropic response of only *MPK3* overexpression lines (n = 10) **(A, B, D)**. Extenal NAA supplementation restored gravitropic response of the *MPK3* overexpression lines (n = 10) **(E-G)**.

### Disruption of *PIN1A* gene dampens gravitropic response by affecting auxin distribution and altering MPK3 activity

Taking hints from a previous study (Xu et al., 2005), we chose to knockout a member of *PIN* family genes that lowers the gravitropic response of rice. Thus, we knocked the *PIN1A* gene out (Supplementary Figure S4A-S4H). Unsurprisingly, the *PIN1A* knockout lines had dampened gravitropic response (Figure 3A), more tiller angle (Figure 3B, C) and decreased plant height (Figure 3D). Knocking the *PIN1A* out also resulted in lowering of auxin level in rice leaves (Figure 3E) due to disruption in auxin transportation. However, surprisingly we found MPK3 activity got enhanced in these lines (Figure 3F) which indicated disruption of PIN1A mediated auxin flow could provide feedback for increasing MPK3 activity in the cells (which in turn contributes to slower auxin transportation by downregulating PINs including PIN1A). Thus, a positive feedback loop involving MPK3 activity and PIN1A-mediated auxin transportation operates in plant cells that regulate gravitropic response or tiller angle in rice.

**Figure 3.**
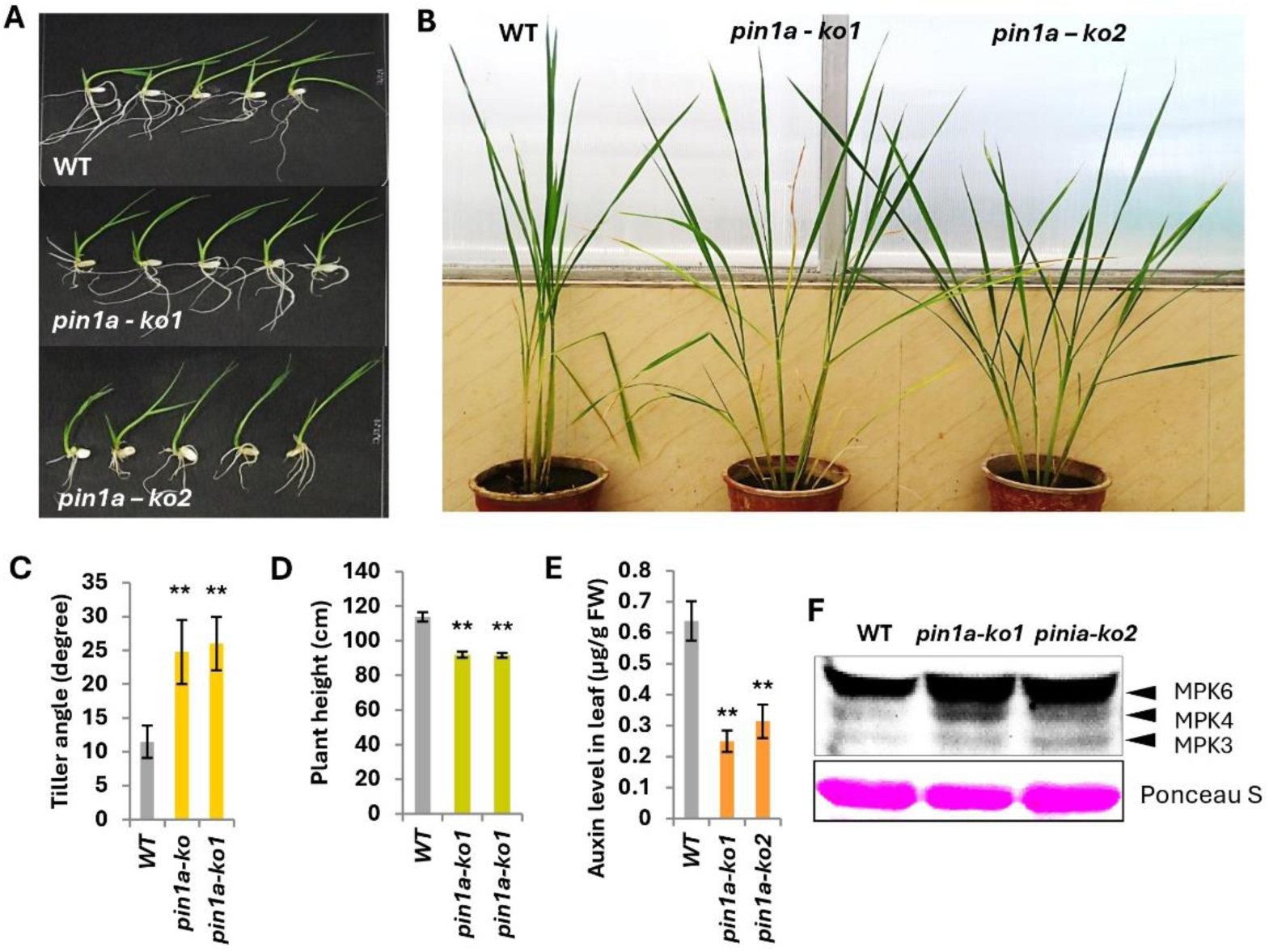
The morphology and physiology of *pin1a* knockout lines. The *pin1a* knockout lines (*pin1a-ko1* and *pin1a-ko2*) had dampened gravitropic response **(A)**, wider tiller angle (n = 8) **(B, C)**, reduced plant height (n = 8) **(D)**, lower auxin content (n = 11) **(E)** and higher MPK3 activity **(F)**.

### MPK3 phosphorylates PIN1A

We wanted to investigate how MPK3 influences the functionality of PIN1A. A previous study in Arabidopsis showed that MPK6 phosphorylates the rice PIN1A homolog (PIN1 in Arabidopsis) and affects its localization behavior (Jia et al., 2026). In the present study, we found MPK3 to physically interact with PIN1A in both *in vitro* (Figure 4A) and *in vivo* (Figure 4D) situations as revealed by pull down and BiFC (Bimolecular Fluorescence Complementation) assays respectively. Further, *in vitro* kinase assay revealed that MPK3 phosphorylates PIN1A (Figure 4I). Analysis of the rice PIN1A sequence revealed presence of three ‘TP’ and two ‘SP’ motifs (the potential MPK phosphorylation sites; Supplementary Figure S5A) in the HL (hydrophilic loop) domain of PIN1A (Supplementary Figure S5B). Among these, all the ‘TP’ and ‘S^351^P’ motifs were found to be hundred percent conserved in all the PIN1A proteins analyzed (Supplementary Figure S5C, S5D). The ‘TP’ motifs of PIN1 are quite well conserved throughout the plant species and are the well-known targets of MPK mediated phosphorylation, while the ‘SP’ motifs have usually been found to be hotspots for PID Kinase mediated phosphorylation (Lanassa Bassukas et al., 2022). However, the study by Jia et al. (2016) revealed S^337^ to be the phosphorylation target of MPK6 in Arabidopsis. The corresponding S residue of Arabidopsis S^337^ in rice was followed by A residue (and hence not a phosphorylation target of MPK3) and another S residue (i.e. S^351^) was found to be ubiquitously conserved. In the present study, thus, we chose to mutate this S^351^ residue into A (for developing the phospho-null variant) and D (for developing phospho-mimic variant) respectively. These mutations did not affect the *in vitro* (Figure 4B, C) and *in vivo* interaction (Figure 4E-H) between MPK3 and PIN1A. However, there was a drastic reduction in phosphorylation signal when S^351^ was mutated to A^351^ (Figure 4J; Supplementary Figure S6). The study revealed S^351^ to be a conserved and a phosphorylation target of MPK3 in rice.

**Figure 4.**
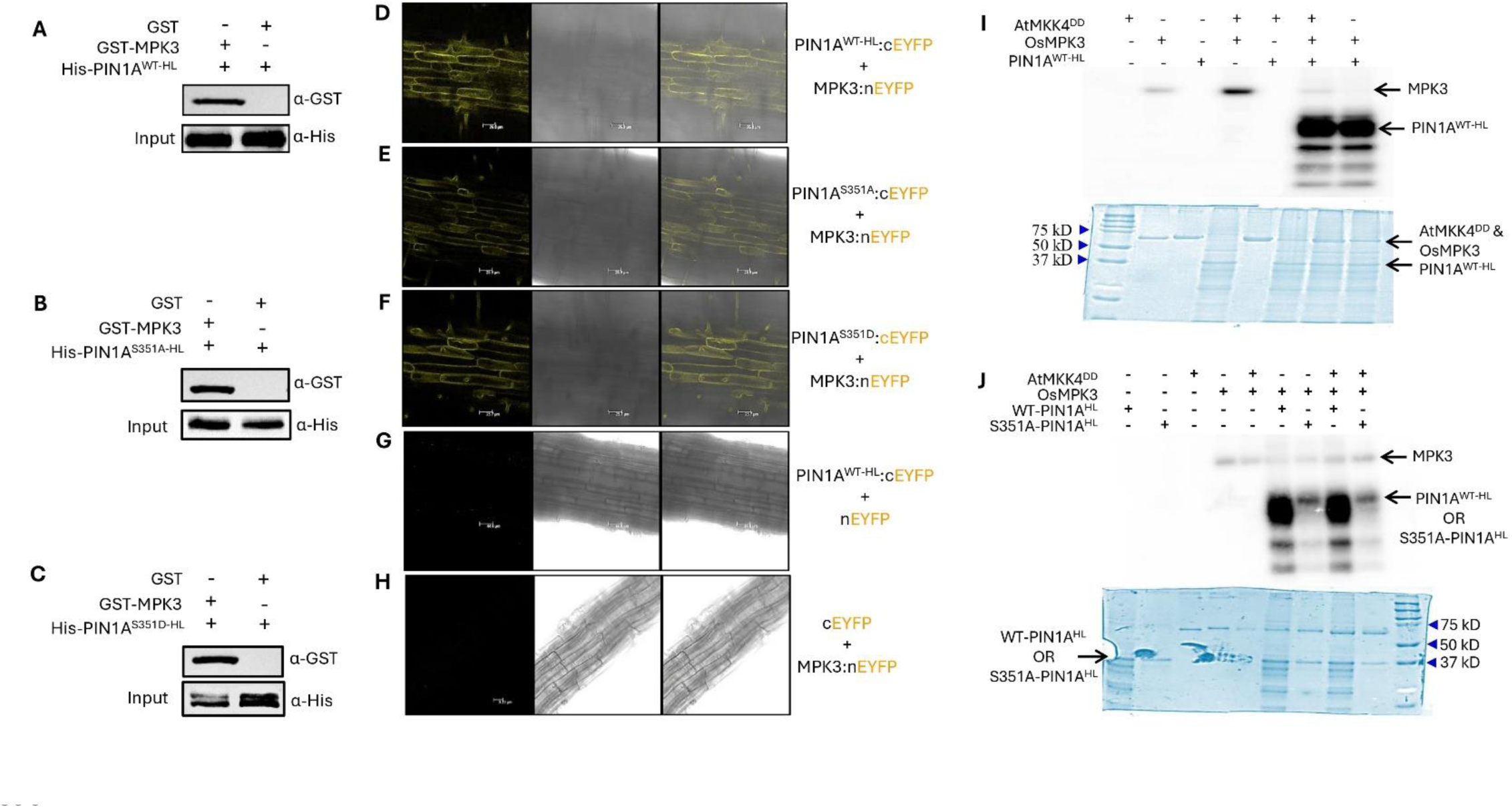
MPK3 interacts with PIN1A and phosphorylates it. The hydrophilic loop (HL) domain of wild-type PIN1A (PIN1A^WT-HL^), phospho-dead PIN1A (PIN1A^S351A-HL^) and phospho-mimic PIN1A (PIN1A^S351D-HL^) interacts with MPK3 both under *in vitro* condition **(A-C)** as determined by *in vitro* pull-down assay as well as under *in vivo* condition as determined by BiFC assay in rice root tissue **(D-H)**. Further, *in gel* kinase assay showed that MPK3 phosphorylates wild-type PIN1A with strong phosphorylation signal **(I)** which got drastically reduced when MPK3 was incubated with phospho-dead PIN1A **(J)**.

### Phosphorylation of PIN1A is vital for endosome mediated trafficking

What is the functional significance of PIN1A phosphorylation? Similar to Arabidopsis, does phosphorylation affect PIN1A localization? If so, how? Curious to find out the answers, we transiently transformed the rice roots and calli with RFP-tagged hydrophilic loop (HL) domain of wild type (WT) or phospho-null (SA) or phospho-mimic (SD) variants of PIN1A. In our previous study (Manna et al., 2024b), we have shown the HL domain of rice PIN1A alone to be enough, for efficiently being targeted to the cell membrane of both root as well as callus cells. The present study revealed that transient transformation of rice tissues with the aforesaid constructs not only illuminated cell membranes with RFP signals, but the signal was also evident in tiny round-shaped vesicles (Figure 5A, B). Surprisingly, the number of vesicles was the highest in case of *PIN1A^SD-HL^* construct followed by *PIN1A^WT-HL^* and *PIN1A^SA-HL^* constructs (Figure 5C, D). To reveal the identity of these vesicles, we transiently co-transformed the rice calli with ER-RFP marker and BiFC constructs of *MPK3* and *PIN1A^WT-HL^*. We found the BiFC signals to co-localize ER signals (Supplementary Figure S7). Thus, these round vesicles were identified to be the ER transport vesicles or endosomes. Further, the phenotype of the PIN1A^HL^-RFP localization signals was similar to the ER localization signal (Supplementary Figure S8A-S8D). Since transport vesicle formation was the highest in the case of *PIN1A^SD-HL^* and the lowest in case of *PIN1A^SA-HL^* construct, it was understood that MPK3-mediated phosphorylation of PIN1A is responsible for endosome-mediated PIN1A trafficking to the cell membrane where masking a vital phosphorylation site lowered transport vesicle formation while mimicking constitutive phosphorylation status increased transport vesicle formation. We also generated the stable transgenic lines of these localization constructs and found that in case of *PIN1A^WT-HL^* and *PIN1A^SA-HL^*constructs, the evidence of transport vesicle formation was not apparent at a given time point (Figure 5E, F) while for the *PIN1A^SD-HL^* construct, there was continuing activity of transport vesicle formation at given time point (Figure 5G, H). In depth observation of the *PIN1A^SD-HL^* stable overexpression line revealed the emergence/coalesce of transport vesicles from/to the cell membrane (Supplementary Figure S9A, S9B), and they had a similar appearance as ER marker (Supplementary Figure S9C). Thus, we found that phosphorylation of PIN1A is vital for its endosome mediated trafficking.

**Figure 5.**
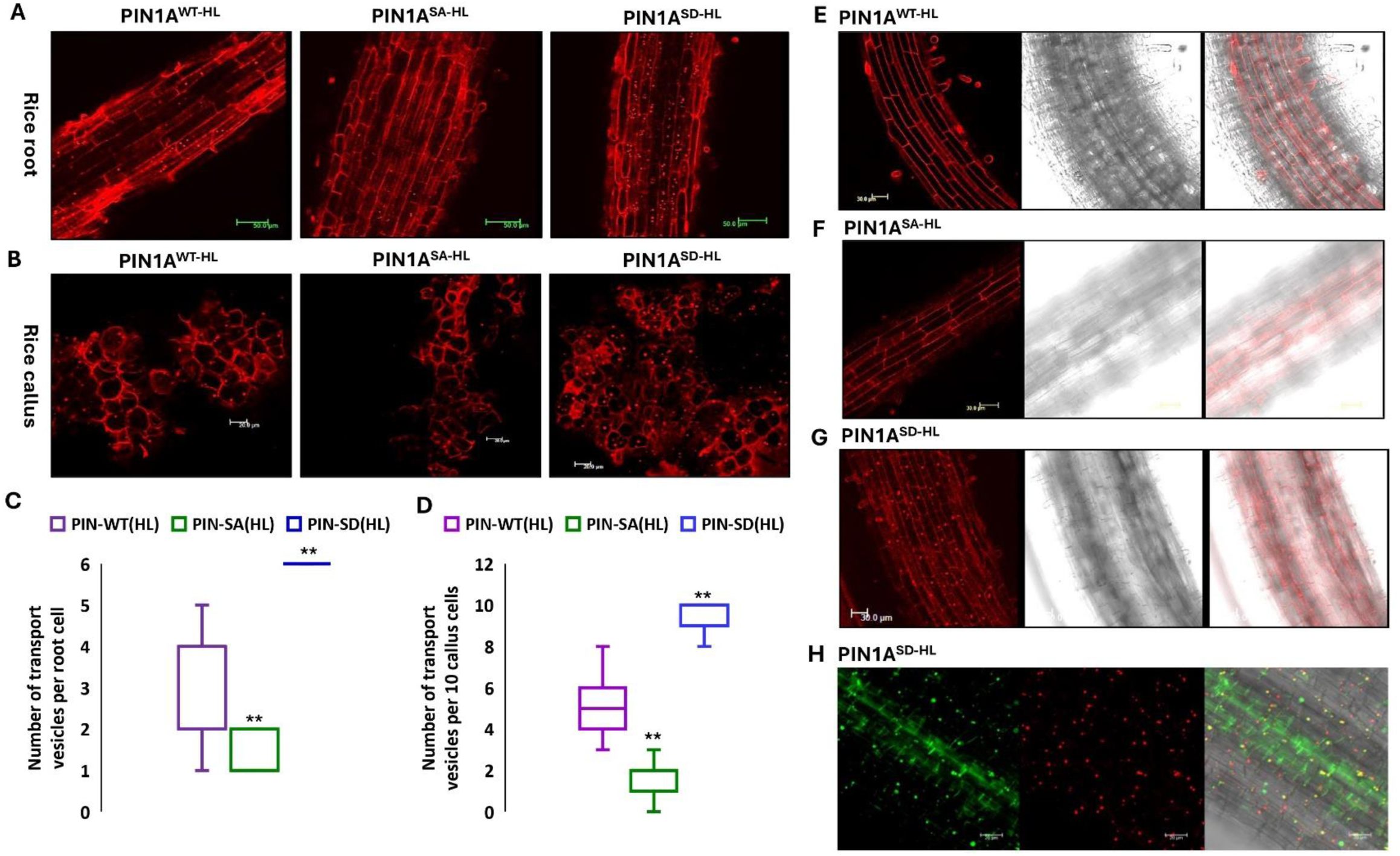
Phosphorylation of PIN1A HL domain facilitates its endosomal trafficking. Transient transformation of rice roots with wild-type PIN1A (PIN1A^WT-HL^), phospho-dead PIN1A (PIN1A^SA-HL^) and phospho-mimic PIN1A (PIN1A^SD-HL^) constructs revealed the highest number endosomal transport vesicle formation in case of PIN1A^SD-HL^ construct and the lowest in case of PIN1A^SA-HL^ construct (n = 15) **(A, C)**. Similar was the situation when rice calli were transiently transformed with these constructs (n = 15) **(B, D)**. While at a given time point endosomal PIN1A trafficking was not evident in the roots of stable transgenics of PIN1A^WT-HL^ and PIN1A^SA-HL^ plants, there was hyperactivity of endosomal PIN1A trafficking in the stable transgenics of PIN1A^SD-HL^ roots **(E-G)**. Counterstaining the PIN1A^SD-HL^ roots with DAPI (here blue colour of DAPI staining has been given a false green colour for better contrast) reveled only some RFP signals to merge with DAPI signals **(H)** as RER (rough ER) signals are also found in nuclei.

To find out if there is any formation of PIN1A transport vesicles when a plant organ moves with continuous change in the direction of movement, we subjected root bits from stable *PIN1A^SA-HL^* transgenics to a continuous rotation motion for four hours following which formation of endosomal transport vesicles was noted. The root bits kept still had very less number of transport vesicle formation while the ones kept under continuous rotation had more transport vesicle formation (Supplementary Figure S10A-S10C). This indicated that PIN1A trafficking takes place during organ moment necessary for differential auxin distribution.

Further, we found that mature *PIN1A^HL-RFP^* overexpression lines had greater tiller angle (Supplementary Figure S11A, S11B), lesser plant height (Supplementary Figure S11C) and lower leaf auxin content (Supplementary Figure S11D) which indicated localization of PIN1A^HL-RFP^ proteins (Supplementary Figure S11E) in the cell membrane would have competitively inhibited localization of endogenous PIN1A transporters (Supplementary Figure S11F) in the cell membrane, the latter having capacity to form desired pore architecture in the cell membrane for auxin efflux, and not the HL domain which alone once integrate in the cell membrane might form abnormal passage for auxin efflux.

### Lesser the PIN1A trafficking, slower the auxin distribution, higher the MPK3 activity and greater the tiller angle

In order to understand physiological implications of PIN1A trafficking, young seedlings of a *pin1a* knockout line were transiently-transformed with either of the empty vector/ *PIN1A^WT^*/ *PIN1A^SA^*/ *PIN1A^SD^* overexpression constructs and their gravitropic response were analyzed. It was observed that the empty vector construct and *PIN1A^SA^* construct couldn’t improve shoot curvature of the *pin1a* knockout seedlings even while effectively transforming them and increasing transcript abundance of the *PIN1A* gene, while the other two constructs could (Supplementary Figure S12A-S12D). This gave a hint that slower rate of PIN1A trafficking (as in the case of *PIN1A^SA^*) is not effective in restoring the normal gravitropic response of the rice seedlings. To check the hypothesis, we generated the stable transgenic lines of *PIN1A^WT^*, *PIN1A^SA^* and *PIN1A^SD^* constructs (their response to external NAA/TIBA are represented in a detailed manner in Supplementary Figure S13). Gravitropic response analysis revealed that *PIN1A^SA^* overexpression line had a slower rate of shoot curvature (Figure 6A, B) and consequently, wider tiller angle (Figure 6C-E) in comparison to the WT, *PIN1A^WT^* and *PIN1A^SD^* overexpression lines. Further, the mature *PIN1A^SA^* overexpression plants were shorter in comparison to the WT and other overexpression lines (Figure 6F). Further, *PIN1A^SA^* overexpression plants had lower auxin content while *PIN1A^WT^* and *PIN1A^SD^* overexpression lines had higher auxin levels in leaf tissue in comparison to WT plants (Figure 6G). Surprisingly, the *PIN1A^SA^* overexpression line was found to have greater MPK3 activity in relation to the rests (Figure 6H) which indicated that lesser rate of PIN1A trafficking caused slower rate of auxin distribution which in turn provided a positive feedback signal to enhance MPK3 activity in the plant cells which we have already shown to be responsible for dampening gravitropic response of rice resulting in its wider tiller angle. Molecular analysis of the transgenic lines showed that all of them had greater *PIN1A* transcript abundance compared to the WT plants (Figure 6I).

**Figure 6.**
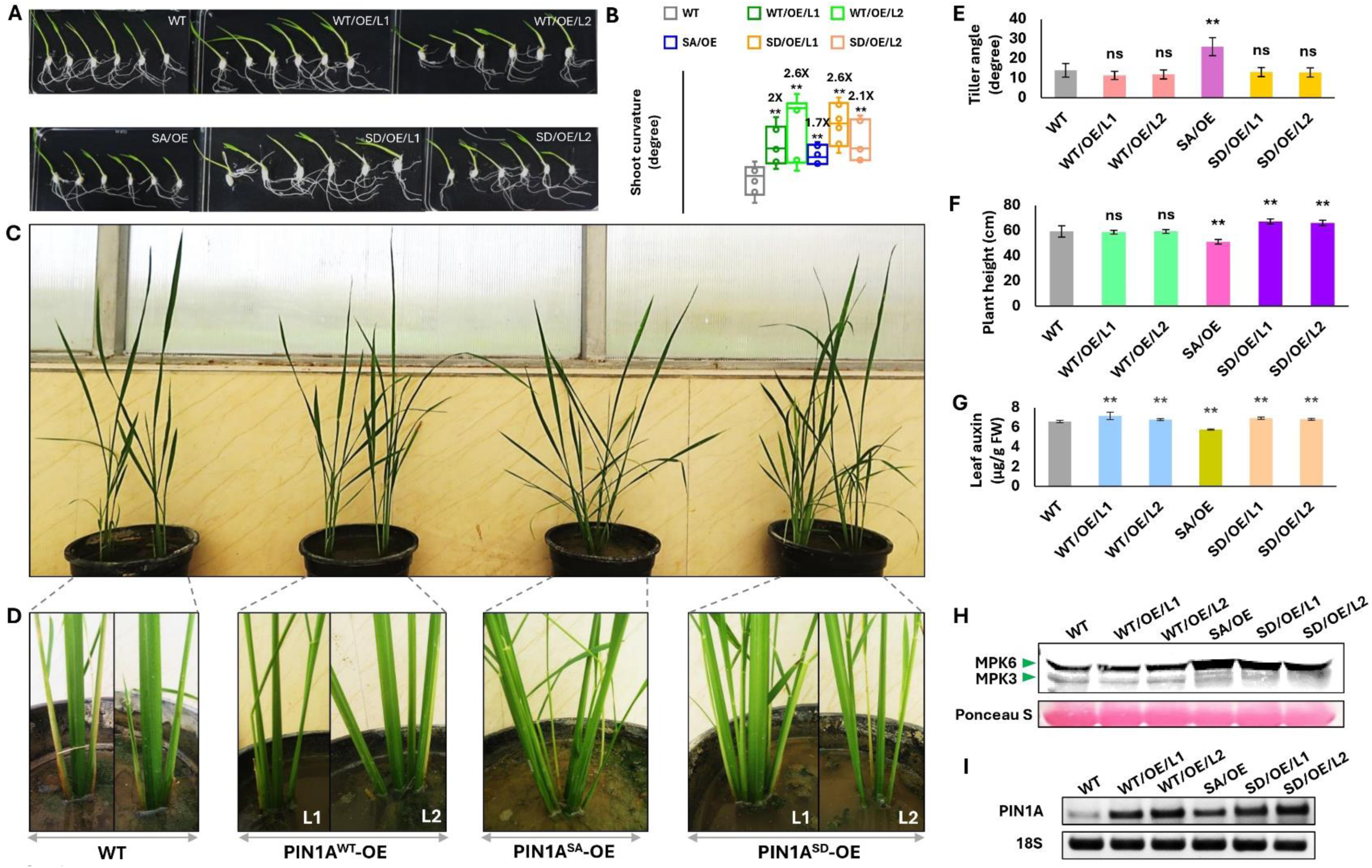
The morphology and physiology of overexpression lines of wild-type *PIN1A* and its variants. Overexpression line of phospho-dead *PIN1A* (SA-OE) had lower gravitropic response (the values over bars designate times more shoot curvature in relation to the wild-type (WT) plants; n = 6) **(A, B)**, greater tiller angle (n = 10) **(C-E)**, lesser plant height (n = 10) **(F)** lower auxin content (n = 8) **(G)** and higher MPK3 activity **(H)** in comparison to the overexpression lines (designated as L1 and L2) of wild-type *PIN1A* (WT-OE) and phospho-mimic *PIN1A* (SD-OE). The expression level of *PIN1A* was more in all the overexpression lines as compared to the WT plants **(I)**.

### Faster auxin transportation delays wound induced tissue senescence in rice

Canher et al. (2020) demonstrated that vascular cell death obstructs polar auxin transport resulting in accumulation of auxin around the periphery cell death area which in turn aids the endodermal cells to undergo periclinal cell division to repopulate the vascular stem cell pool. This prompted us to enquire if PIN1A mediated polar auxin flux could affect wound-induced senescence in rice. To understand this, the *DR5-gus* expressing rice leaf bits were pricked with a needle and left at room temperature for 24 hours either in water or in water containing 1 mg/L NAA. The gus staining around the periphery of tissue injury showed accumulation of auxin (Figure 7A), while in the case of NAA treatment, gus staining was also evident beyond the injury periphery indicating auxin flux into or out of the injured area (Figure 7B). Tissue injury triggers senescence and leaf yellowing in plants (Lim and Nam, 2005). Measurement of the extent of leaf yellowing following tissue injury revealed that yellowing was significantly less when the leaf bits were incubated in the presence of auxin (Figure 7C, D). Auxin treatment was also found to upregulate the expression of some PIN members including *PIN1A*, *PIN1B*, *PIN1C* and *PIN5C* (Figure 7E). Further, leaf auxin content was also found to increase after tissue injury and also when the injured leaves were incubated in the presence of NAA (Figure 7G). Wounding also resulted in MPK (MPK3 and MPK6) activation while NAA treatment suppressed wounding mediated activation of MPKs (Figure 7F). Analysis of leaf senescence or leaf yellowing (following pricking the leaves by needle) of the *MPK3* overexpressing and knockout lines revealed that leaf yellowing was faster in the overexpression lines than WT while leaf yellowing was significantly slower in the knockout line (Supplementary Figure S14A, S14B). The experiment thus revealed that an increase in auxin flux delays wounding mediated tissue senescence by lowering activation of MPK3.

**Figure 7.**
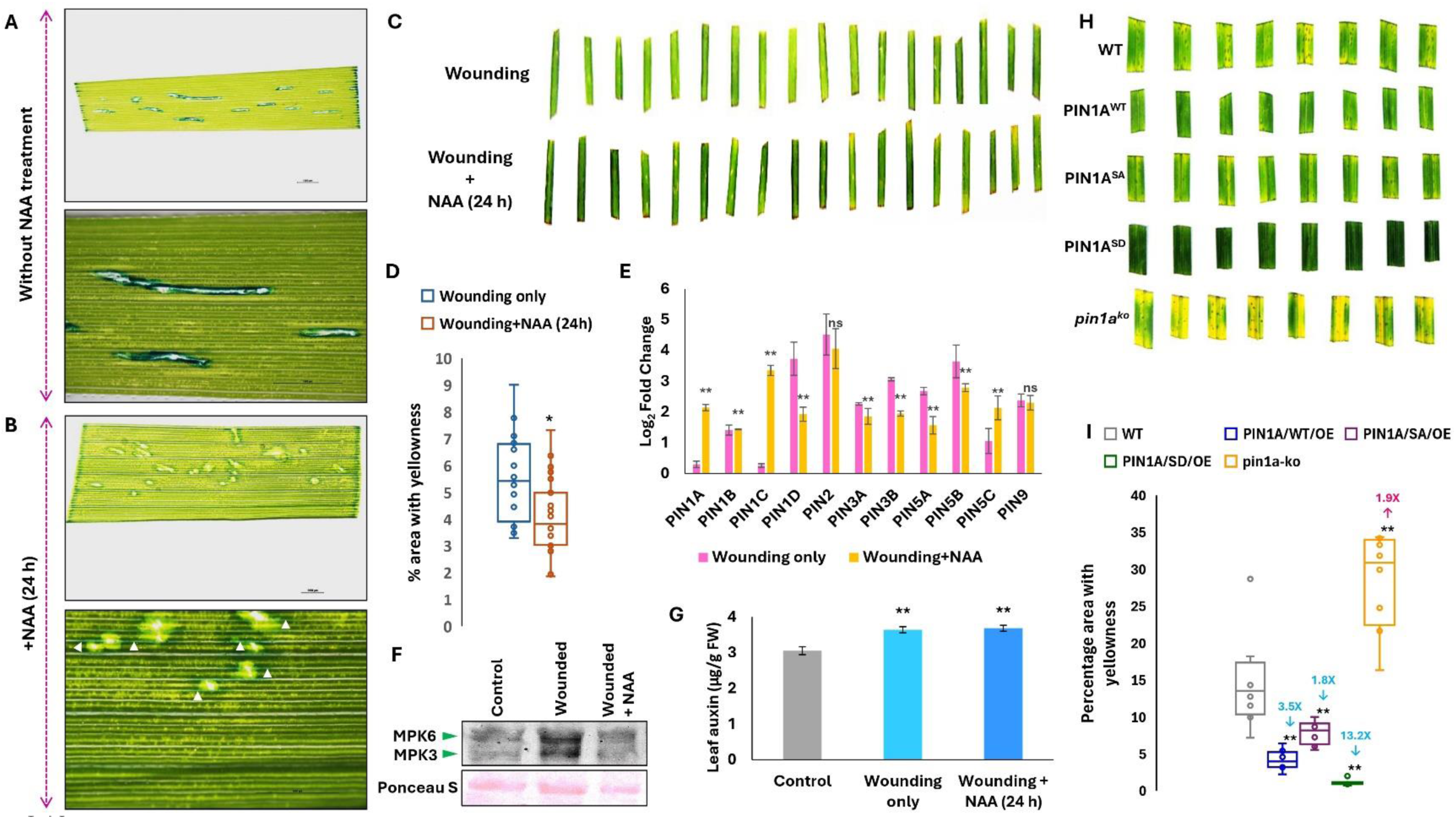
Increase in auxin flux delays tissue senescence in rice. **(A)** There is increase in auxin accumulation around periphery of wounds as revealed by gus staining of the *DR5-gus* expressing rice leaves. Auxin (NAA) treatment not only increased auxin flux **(B)** but also lowered yellowing (or tissue senescence) following mechanical wounding (n = 23) **(C, D)**. **(E)** Wounding alone or in combination with NAA treatment upregulated *PIN* genes’ expression depicting upsurge of auxin transportation happens following tissue injury (n = 3). NAA treatment however significantly increased the expression of *PIN1A*, *PIN1C* and *PIN5C* post wounding. **(F)** While wounding alone increased MPK3 activity, combined wounding and NAA treatment suppressed MPK3 activity in rice leaves. **(G)** A given piece of rice leaf had more auxin content following wounding alone or in combination with NAA treatment depicting increase in auxin flux happens post wounding (n = 4). **(H, I)** Analysis of effect of wounding on leaves of WT and various *PIN1A* overexpression/ knockout lines (line number 1 was used in all the cases) revealed rapid tissue senescence in *pin1a* knockout line (*pin1a^ko^*) followed by wild-type (WT), phospho-dead PIN1A overexpression line (*PIN1A^SA^*), wild-type PIN1A overexpression line (*PIN1^WT^*) and phospho-mimic PIN1A overexpression line (*PIN1A^SD^*) (n = 10).

Further, analysis of the leaf yellowing pattern in the *PIN1A* overexpression and knockout lines revealed that *PIN1A^SD^* leaves retained maximum greenness followed by *PIN1A^WT^*, *PIN1A^SA^* and WT leaves, while *pin1a* knockout leaves displayed rapid senescence (Figure 7H, I). Thus, it was indeed apparent that the faster the auxin flux, slower the tissue senescence. Additionally, there was greater endosomal PIN1A trafficking following tissue injury which again emphasized heightening of auxin flux following tissue damage (Supplementary Figure S15A-S15C).

### PIN1A expression and trafficking regulate wounding responses in rice

In order to see the influence of wounding on overall plant physiology, MPK3 activity and auxin accumulation, leaves of WT, *PIN1A^WT^*, *PIN1A^SD^* overexpression and *pin1a* knockout plants were scratched with sandpaper and left for three days in the greenhouse with normal irrigation. The *PIN1A^WT^* and *PIN1A^SD^* plants had milder necrotic lesions, while *pin1a* knockout plants had greater and more prominent necrotic lesions as compared to the WT plants (Figure 8A-D). There was a clear induction of MPK3 activity in WT following tissue injury while enhancement of MPK3 activity was not prominent in all the *PIN1A* overexpression and knockout lines following tissue injury (Figure 8E). However, WT and all the *PIN1A* lines had increased auxin accumulation upon tissue injury (Figure 8F-I). To assess the effect of wounding on the physiology of the rice plants, we measured the photosynthetic parameters of the plants under control conditions and following tissue injury. Under control conditions, the photosynthesis rate of *PIN1A^WT^* and *PIN1A^SD^* plants was higher than WT, while in the case of *pin1a* knockout plants, the photosynthesis rate was significantly lower than the rest. Tissue injury led to a lowering of the photosynthesis rate in all the plants, while maintaining a trend similar to the one seen under the control condition (Figure 8J). Stomatal conductance was higher in *PIN1A* overexpression lines while the knockout line had lower stomatal conductance than the WT plants under control condition. Tissue injury lowered stomatal conductance in all the plants, and it was lower in both the overexpression and knockout lines than the WT plants following tissue injury (Figure 8K). Under control condition, intercellular CO_2_ concentration was higher in *PIN1A^WT^* and *pin1a* knockout plants while it was higher in case of *PIN1A^SD^* compared to the WT plants. However, upon tissue injury, intercellular CO_2_ concentration of the *PIN1A^WT^* and *PIN1A^SD^* plants got significantly reduced (Figure 8L). The transpiration rate was higher *in PIN1A^WT^* and *PIN1A^SD^* plants and lower in *pin1a* knockout plants than the WT under normal conditions, while after wounding, transpiration rate was significantly lower in the *PIN1A^WT^* and *PIN1A^SD^*compared to the WT plants (Figure 8M). Under control condition, water use efficiency was significantly more in the *PIN1A^SD^* plants while in the wounded plants, water use efficiency was significantly higher in the *PIN1A^WT^* and *PIN1A^SD^* plants in relation to the wounded WT plants (Figure 8N). Overall, the results indicated that *PIN1A^WT^* and *PIN1A^SD^* plants sustained better physiology in the form of maintaining higher photosynthesis rate and water use efficiency while lowering the rate of transpiration than the WT plants following tissue injury.

**Figure 8.**
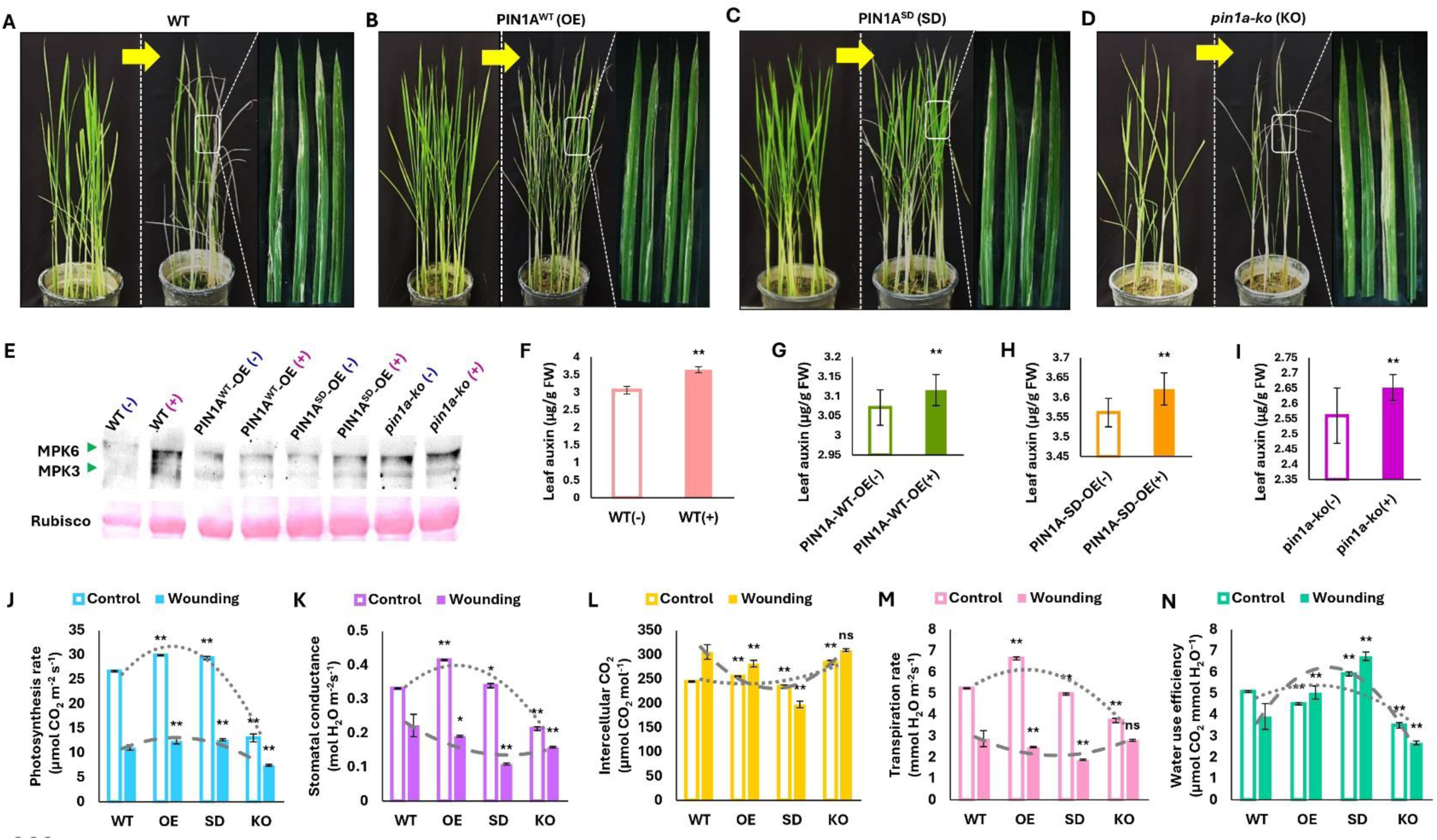
The effect of mechanical wounding on various *PIN1A* lines. **(A-D)** Mechanical wounding of intact rice leaves by rubbing with a sandpaper caused visibly more tissue damage in *pin1a* knockout line (*pin1a-ko*) and wild-type plants (WT) than the wild-type PIN1A overexpression line (*PIN1^WT^*) and phospho-mimic PIN1A overexpression line (*PIN1A^SD^*). **(E)** The pattern of MPK3 activation before and after tissue injury. **(F-I)** Graphs showing increase in auxin content of rice leaves in WT and overexpression/ knockout lines post tissue injury (n = 23). Various photosynthetic parameters namely photosynthesis rate **(J)**, stomatal conductance **(K)**, intercellular CO_2_ level **(L)**, transpiration rate **(M)** and water use efficiency **(N)** of aforesaid rice plants before and after wounding (n = 5). The dotted curves over each graph are polynomial curves representing the trend of change in photosynthetic parameters among the plant types.

Further, the mature WT and various *PIN1A* lines growing in the net house were found to naturally contract brown leaf spot disease (the fungal pathogen, *Biploaris oryzae* responsible for the disease was subsequently isolated and characterized by visualization of characteristic spore formation, molecular confirmation by pathogen gene specific PCR and observing emergence of disease symptoms after inoculation the fungal spores into WT and various *PIN1A* lines; Supplementary Figure S16A-S16F) and closer observation of the leaves revealed a difference in the number and size of necrotic spots among the WT and *PIN1A* lines. Among many factors, pathogen-mediated tissue injury is an important cause of the development of necrotic lesions (Vidhyasekaran et al., 1992). In relation to the WT plants, *PIN1A^WT^*, *PIN1A^SA^* and *PIN1A^SD^* lines had smaller sized lesions while lesion lengths were longer in the case of *pin1a* knockout lines (Supplementary Figure S17A, S17B). Furthermore, analysis of the number of lesions per 10 cm length revealed that in comparison to the WT plants, *PIN1A^WT^* and *PIN1A^SD^* lines had a lesser number of lesions while the *pin1a* knockout lines had a greater number of disease spots (Supplementary Figure S17A, S17C). This indicated PIN1A-mediated enhanced auxin circulation can also counter pathogen-mediated wounding in rice.

## Discussion

### MPK3 mediated phosphorylation of HL-domain of PIN1A regulates its trafficking

The directional transportation of auxin is required throughout a plant’s life cycle as the role of auxin is critical for all aspects of plant physiology and development including but not limited to embryogenesis (Karami et al., 2023), tissue differentiation (Su et al., 2011), organogenesis (Suzuki et al., 2023), trophic growth (Barbosa and Schwechheimer, 2014) and tissue regeneration (Canher et al., 2020; Hoermayer et al., 2020). The directionality of auxin flow or its polar transportation is achieved to a large extent by PIN-FORMED (PIN) auxin efflux transporters (Lanassa Bassukas et al., 2022; Luschnig and Friml, 2024). Phosphorylation of PIN proteins is known to facilitate PIN trafficking and their polar localization in the cell membrane (Zourelidou et al., 2014; Zwiewka et al., 2019). So far, four classes of protein kinases are known to phospho-regulate PIN proteins at their HL domain: (i) In the AGCVIII family of serine/threonine kinases (Dhonukshe et al., 2010; Grones et al., 2028; Lanassa Bassukas et al., 2022), (ii) the plasma membrane-localized CRKs (Ca^2+^/CALMODULINDEPENDENT PROTEIN KINASE-RELATED KINASEs) (Baba et al., 2019a; 2019b; Rigó et al., 2013), (iii) the auxin-regulated receptor CAMEL (CANALIZATION-RELATED AUXIN-REGULATED MALECTIN-TYPE RECEPTOR KINASE) (Hajný et al., 2020) and (iv) the MPKs which are well known to phosphorylate three highly conserved T residues (i.e. T^227^, T^248^ and T^286^) located within three TPRXS motifs in Arabidopsis (Dory et al., 2018). A study by Jia et al. (2016) discovered that S residue (S^337^) can also be a target of MPK-mediated phosphorylation that has a potential to regulate PIN1 polarity in Arabidopsis (Zhang et al., 2010). In the present study, we discovered a highly conserved S residue (S^351^) in rice PIN1A which is a phosphorylation target of MPK3 (Figure 4). The mutation of the S residue into phospho-null variant (i.e. A) suppressed endosome mediated PIN1A trafficking while its mutation into phospho-mimic form (i.e. D) resulted in more PIN1A trafficking activity inside the cell (Figure 5). The study by Dory et al. (2017) witnessed that conversion of T^227^, T^248^ and T^286^ residues of PIN1 to phospho-mimic residues (i.e. E) resulted in the formation of aggregates inside cells that corresponded well with the ER marker, and they hypothesized that formation of these aggregates might be responsible for interference in PIN1 trafficking in Arabidopsis. We also witnessed the formation of such aggregate bodies when S^351^ of PIN1A was mutated to phospho-mimic form (i.e. D^351^) which were found to be ER vesicles or endosomes (Supplementary Figure S7, S8, S9). But rather than interfering with PIN1A trafficking, they aided the process, as higher PIN1A trafficking activity in the case of PIN1A^S351D^ increased auxin flux in rice as compared to the WT plants while in the case of PIN1A^S351A^, PIN1A trafficking activity got significantly reduced casing reduction in auxin flux and dampening of shoot gravitropism in rice.

### Gravistimulation and organ movement trigger endosome-mediated PIN1A trafficking thus regulating the pace of auxin distribution and shoot gravitropism

During gravistimulation, the shoot starts bending upward displaying negative geotropism property (Figure 1, 2, 3, 6). Thus, plants can sense gravity. The classical starch–statolith hypothesis postulated that specialized cells containing amyloplasts can sense gravity where their sedimentation enables plants to sense gravity (Fujihira et al., 2020; Fukaki et al., 1998; Morita et al., 2024; Sack, 1997). Gravity sensing triggers movement in plant organs. In the present study, we showed that movement in plant organ (here, root tissue) increased the incidence of endosome mediated PIN1A trafficking (Supplementary Figure S10). Ditengou et al. (2008) demonstrated that mechanical bending of a root portion in Arabidopsis stimulated redistribution of plasma membrane-localized PIN1 that preceded auxin dependent gene transcription for initiation of lateral root primordium from the convex side of the bent. We also witnessed that mechanical stimulation of organ movement increases the PIN1A trafficking in rice (Supplementary Figure S10). Increase in PIN1A trafficking is expected to increase auxin flux and in line with this, we witnessed increase in leaf auxin content in the *PIN1A^WT^*and *PIN1A^SD^* overexpression lines which further led to significant increase in their shoot gravitropic response in comparison to the WT and *PIN1A^SA^* seedlings (Figure 6). The classical Cholodny–Went hypothesized that asymmetric auxin redistribution causes differential growth on lower and upper side of an organ that enables plants to bend, take a curvature (that is, display tropisms) (Wang et al., 2022). In the present study, we also showed greater auxin accumulation in the convex side of the root curvatures which not only led to more cell expansion in the convex side which facilitated curvatures in the rice roots (Figure 1). Taken together, the present study shows that gravistimulation mediated organ movement is one of the leading causes of PIN1A trafficking and MPK3 mediated phosphorylation of its S^351^ residue aids the process. This helps in regulating gravitropic response and hence tiller angle in rice. It is also important to mention here that shoot gravitropism greatly influences tiller angle in rice (Harmoko et al., 2016; Huang et al, 2021; Hu et al., 2020; Zhang et al., 2018) and the same is also evident from the present study (Figure 3, 6).

### Wounding also enhances endosome mediated PIN1A trafficking that determines pace and extent of tissue senescence

Plants face many types of tissue damage, including the ones caused by herbivory and other forms of physical wounding like breakage of plant organ because of wind, trampling by animals or pathogen mediated cell damage (Savatin et al., 2014; Vega-Muñoz et al., 2020). The sessile plants have evolved various ways to regenerate tissues following tissue injury and wounding. Whatever be the multiple complex molecular signaling pathways behind tissue regeneration, they ultimately culminate to auxin accumulation around periphery of damaged tissue (as death of vascular stem cells death results in obstruction of auxin flux) which in turn enables endodermal cells to undergo cell expansion and periclinal cell division for re-populating the vascular stem cell pool that heals the wounds (Canher et al., 2020; Hoermayer et al., 2020). We found an incidence of higher PIN1A trafficking following tissue damage (Supplementary Figure S15). Additionally, we also witnessed that increase in PIN1A trafficking increased auxin flux inside the plants as evidenced by higher auxin content in the *PIN1A^WT^* and *PIN1A^SD^* plants in comparison to WT, *PIN1A^SA^* and *pin1a* knockout plants (Figure 3, 6). The greater auxin flux would have facilitated faster auxin built up around the periphery of damaged tissue thus slowing up tissue injury mediated leaf senescence in rice.

Pathogens also damage plant cells by producing enzymes that injure healthy tissues (Hoermayer et al., 2020). Necrotrophic fungi like *Bipolaris oryzae* (causes brown spot disease in rice) also cause wounding in rice tissue by producing a toxin that suppresses phenol metabolism in rice leaves (Vidhyasekaran et al., 1992). In leaves of grown-up rice plants, the disease symptoms appear as dark brown oval necrotic spots all over the leaves which coalesce to form bigger necrotic lesions or drying up of leaves from tips in the susceptible plants (Supplementary Figure S16, S17). Increased PIN1A trafficking and consequent increase in auxin flux in the *PIN1A^WT^* and *PIN1A^SD^* plants was found to reduce the incidence of development of necrotic lesions following pathogen attack (Supplementary Figure 17). Earlier Sun et al. (2018) found that increased expression of endogenous *PIN1A* led to conferring resistance to sheath blight disease (caused by *Rhizoctonia solani*, a necrotrophic fungus) in rice. Further, Chu et al. (2021) also showed that upregulation of *PIN1A* provides resistance to sheath blight disease in rice. The present study thus revealed the importance of PIN1A trafficking in countering not only tissue senescence following an injury but also providing resistance to a devastating rice pathogen, *Bipolaris oryzae* which caused the infamous Bengal famine that starved about two million people to death during 1942 – 1943 (Bisen et al., 2015; Jatoi et al., 2019).

### Regulation of dual gravitropic and wounding response in rice by a positive feedback loop

We found that when *PIN*s expression is downregulated (during TIBA treatment) or *PIN1A* was knocked out, lower auxin flux led to enhanced MPK3 activation (Figure 1, 3) which in turn caused *PIN* downregulation and lower auxin flux (Supplementary Figure S2). The opposite was the case when auxin content of tissue increased (Figure 1, 6, Supplementary Figure S3). We do not know how MPK3 regulates transcription of *PINs* especially *PIN1A*. We believe that it might happen via phosphorylation mediated degradation of a transcriptional activator of *PIN1A* (such as LPA1; Chu et al., 2021) by MPK3. Further, lower tissue auxin content was associated with dampened gravitropic response (or wider tiller angle) and faster tissue senescence following tissue damage (Figure 1, 3, Supplementary Figure S17) while higher tissue auxin content not only improved gravitropic response of rice but also slowed down tissue senescence and development of necrotic lesions during pathogen infestation (Figure 6, 7, Supplementary Figure S17). Thus, we found that MPK3 activity is regulated by auxin flux which itself is regulated by MPK3 mediated PIN1A phosphorylation and its trafficking. Based on these observations, we propose that a positive feedback loop encompassing ‘MPK3-PIN1A trafficking-auxin flux’ trio regulates dual gravitropic and wounding response in rice (Figure 9).

**Figure 9.**
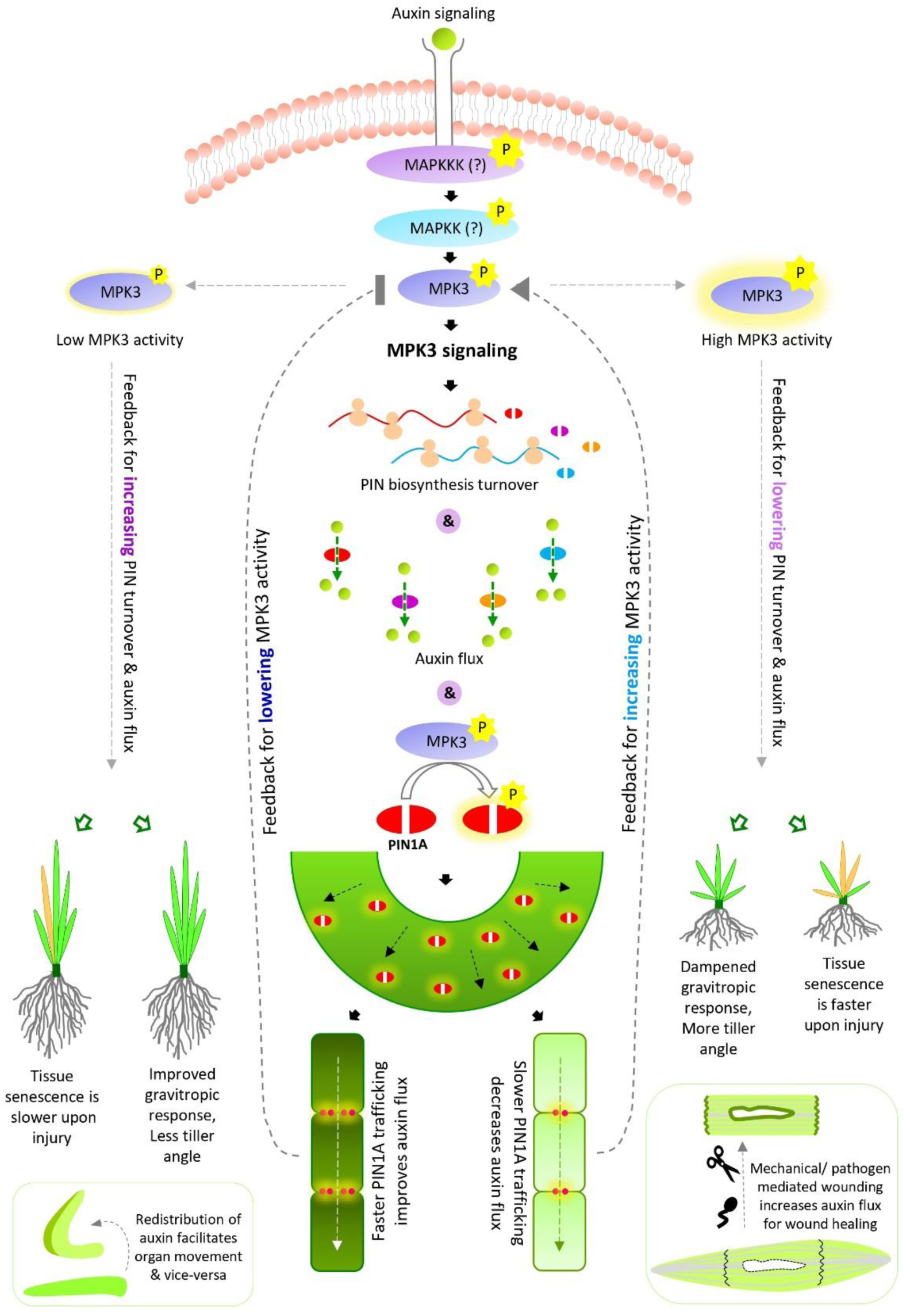
The positive feedback loop encompassing MPK3 activity, PIN1A trafficking and auxin flux regulate dual gravitropic and wounding response in rice. The schematic diagram showing overview of the present research as follows: MPK3 regulates transcription of *PIN* genes including *PIN1A* (probably by regulating the activity of transcription factors that influence the expression of *PIN1A* via their phosphorylation), auxin flux and PIN1A trafficking via its phosphorylation. When PIN1A trafficking is rapid, auxin flux is faster, and this situation provides a positive feedback signal to lower MPK3 activation which in turn leads to upregulation of *PINs* and concomitant increase in auxin flux. Such a positive feedback loop favors improved gravitropic response/ narrow tiller angle and slower senescence following tissue injury. On the other hand, when PIN1A trafficking slows down, there is lesser symplastic flow of auxin which provides a positive feedback signal to activate MPK3 in the cell which in turn downregulates *PIN* genes’ expression resulting in slower auxin flux inside the plant. Such a positive feedback loop dampens gravitropic response of rice thus increasing its tiller angle and it also favors rapid senescence following tissue injury.

## Materials and Methods

### Plant material and growth conditions

Taipei309 variety of *Japonica* rice was the WT (wild type) plant, and all the transgenic and knockout lines were made in this rice variety. The *in vitro* plate level experiments were carried out by germinating and growing the rice seedlings on half strength Murashige and Skoog (MS) basal medium (i.e., 2.2 g/L of MS basal salt; Duchefa, catalogue number: M0221.0050 + 15 g/L sucrose; HiMedia, catalogue number: PCT0607 solidified with 4 g/L phytagel; Sigma, catalogue number: P8169; pH of the media was maintained at 5.8). The 1 µM DEX (Dexamethasone; Sigma, catalog number: D4902; Singh et al., 2019) treatment was given either by adding required amount of DEX to MS medium after autoclaving and cooling the same or by supplementing one-fifth strength MS solution (Manna et al., 2024a) with DEX (Figure 2A, 2B, 2E, Supplementary Figure S2, S14). The NAA (@ 1 mg/L) or TIBA (@ 2 mg/L) treatments were given by supplementing the solid or liquid media with the respective chemicals (Figure 1F, 1K, 2F, 7A-C, Supplementary Figure S13). The seedling level experiments were carried out by growing them in rice tissue culture growth room having a temperature of 27°C, 14 h light/10 h dark photoperiod cycles and 1000 lux light intensity. For seed multiplication and experiments with mature plants (Figure 8), the plants were grown in in pots filled with a 3:1 mixture of soil and cow dung manure in the greenhouse where the growth conditions were similar to growth room with relative humidity maintained at 70–80 % level.

### Construction of plant transformation cassettes and primers used in the study

The *PIN1A* gene was PCR amplified from cDNA of rice seedlings subjected to 3h of 1 mg/L NAA treatment. The making of localization construct of *PIN1A^WT-HL-RFP^*in pGWB454 plant transformation vector is mentioned in detail in Manna et al. (2024b). Similarly, the *PIN1A^SA--HL-RFP^* and *PIN1A^SD--HL-RFP^*localization constructs were made. We used overlap extension PCR (Hilgarth and Lanigan, 2019) to develop the *SA* or *SD* variants of *PIN1A* gene. The various *PIN1A^HL^* genes were cloned in pET-28a vector and the 6X-His tagged proteins were expressed in Arctic cells of *E. coli*. The BiFC (Bi-molecular Fluorescent Complementation) constructs were made in pSITE-nEYFP (ABRC CD3-1648) and pSITE-cEYFP (ABRC CD3-1651) vectors respectively. The overexpression constructs of full length *PIN1A* and its variants were made in pANIC6b vector. The detailed description of making the *PIN1A* knockout construct is mentioned in Manna et al. (2024b) and the knockout construct of rice *MPK3* genes was made in the same way. The primers used in this study are mentioned in Supplementary Table S1.

### Stable transgenic making and transient transformation

The authors of this paper have developed a fail-safe rice transformation protocol for developing stable rice transgenics as well as for carrying out transient transformation in *Japonica* rice background (Manna et al., 2024b). The same protocol was used here to develop stable transgenic lines (Figure 3B, 5E-G, 6A, 6C, Supplementary Figure S11A) as well as for transiently transforming the rice seedlings (Figure 4D-H, 5A, 5B, Supplementary Figure S12) for localization and BiFC studies.

### Microscopy

Stereo microscopic visualization of *DR5-gus* expressing rice root (Figure 1A) and leaves following tissue injury (Figure 7A, B) was done with the help of Leica MZ12 stereomicroscope and all confocal imaging for visualizing roots under UV spectrum (Figure 1B-D) or for detecting the YFP/RFP/DAPI (Figure 4A-H, Figure 5, Supplementary Figure S7-S10, S15) signals was performed by using Leica TCS SP8 confocal microscope at a gain % of 800 V with 70% laser intensity to maintain consistency.

### Measuring gravitropic response of the rice seedlings

For measuring rice seedling’s gravitropic response, the surface sterilized rice seeds were germinated over half strength-MS medium in square shaped petriplates and allowed to grow vertically for 7-10 days during which they attained 3-4 cm length. After that, the plates were inverted by 90° and the shoot curvature angle was measured after 3 days when the horizontally placed seedlings started bending upward due to negative geotropic nature of the shoots.

### RNA isolation and analysis of gene expression

RNA isolation and the semi-qRT or qRT-PCR (quantitative real-time reverse transcriptase polymerase chain reaction) for analyzing gene expression were carried out following the protocol mentioned in Manna et al. (2024b) and Manna et al. (2022).

### Auxin quantification

The authors of this paper developed a simplified and sensitive plant auxin estimation protocol (Manna et al., 2024c). The same protocol was used here for estimating auxin contents of root and shoot tissues.

### *In-vitro* kinase assay

*In-vitro* kinase assay was carried out as per the protocol described by Takahashi et al. (2007). In brief, the proteins (kinase and substrate) were incubated in 10 µL of kinase reaction buffer (50 mM Tris–HCl, pH 7.5, 1 mM DTT, 10 mM MgCl_2_, 10 mM MnCl_2_, 50 mM ATP, and 0.037 MBq of (γ^32^P ATP) [60 Ci/mmol]) at 30 °C for 30 min. Thereafter, the Kinase reactions were by adding 2X SDS loading dye and heating for 5 min at 95 °C to stop the reaction. Reaction products were run on SDS-PAGE gel and signals were detected by phosphor imager (Typhoon, Phosphor Imaging System, GE Health Care, Life Sciences) and presence of proteins were analyzed by staining with Coomassie Brilliant Blue R 250 stain.

### Plant protein extraction and Western Blotting

Crude plant protein extract was isolated from rice root or shoot tissues using kinase extraction buffer (50 mM HEPES (pH 7.5), 5 mM EGTA, 5 mM EDTA, 10 mM dithiothreitol, 10 mM Na_3_VO_4_, 10 mM NaF, 50 mM β-glycerolphosphate, 1 mM phenylmethylsulfonyl fluoride, protease inhibitor cocktail, 10% glycerol). The proteins were quantified with Bradford reagent (Bradford, 1976). Approximately, 30 μg of plant protein was run on 12 % SDS-PAGE and thereafter ran in the western blotting assembly at 25 mA for 1 h for transferring the proteins from gel to membrane. After the transfer was over, the membrane was stained with ponceau stain and washed with water. The membrane was kept in blocking buffer for 16 h at 4 °C and washed thrice with washing buffer (TBST) for 10 minutes each. After washing, the membrane was incubated with primary antibody; pTEpY (dilution 1:15,000) for 1 h. The membrane was again washed thrice for 10 min each with TBST and incubated with secondary antibody (dilution of 1:15,000) in blocking solution for 1h. Membrane was then washed with TBST and Supersignal west Pico chemiluminescent substrate (Pierce, USA) was utilized for detecting the signal.

### Injuring rice leaves and measuring percentage area with yellowness

Even sized leaf bits were cut out from respective WT or *PIN1A* overexpression or knockout lines were pricked 10 times with the help of a syringe needle. The injured leaf bits were then placed over moist filter paper (kept in petriplates) and incubated at 28 °C with lids of petriplates covered for three days before photographing and analyzing. The individual leaf images were analyzed by ImageJ software to determine percentage area with yellowness.

### Photosynthetic parameters

Photosynthesis rate and its related physiological parameters were measured with the help of Li-COR 6400XT portable photosynthesis system (LI-COR, USA) on the fully expanded leaves of 60-days-old rice plants (before and after scratching the leaves uniformly using sandpaper; Figure 8A-D) using the method described by Jonwal et al. (2023).

### Bioinformatic analyses (protein structure prediction, multiple sequence alignment, PIN1A pore diameter profile)

Multiple sequence alignment of PIN proteins was done using the Clustal Omega program (https://www.ebi.ac.uk/jdispatcher/msa/clustalo) with default parameters and the aligned images were further visualized using Jalview software (Supplementary Figure S5C, S5D). The pore morphology of PIN proteins was predicted using PoreWalker software (Supplementary Figure S11G-I) (Pellegrini-Calace et al., 2009). The three-dimensional (3D) structures of proteins were generated using PHYRE2 Protein Fold Recognition Server (https://www.sbg.bio.ic.ac.uk/phyre2/html/page.cgi?id=index) (Supplementary Figure S11E, S11F).

### Identification of fungal pathogen causing leaf spot disease in rice

The scrapings from disease spots were placed in a drop of water over a slide and watched under confocal microscope in bright field to visualize fungal spores if any (Supplementary Figure S16A, S16B). Thereafter, leaf bits of about 1.0 × 0.4 cm dimension containing disease spot were cut from infected rice leaves and surface sterilized with 70 % ethanol for 5 minutes followed by washing thrice with sterile water and blot drying briefly (1 min) before placing over PDA (Potato Dextrose Agar; HiMedia, catalog number: MH096) medium. The PDA plates were then kept in dark inside a 28 °C incubator for five days. Two types of fungal colonies appeared (Supplementary Figure S16C) which were analyzed under confocal microscope to find out sporulation (Supplementary Figure S16D). The genomic DNA was also isolated from these fungal colonies following the protocol described by Conlon et al. (2022). The molecular confirmation of these fungal colonies was done by PCR with *Helminthosporium oryzae*’s *GAPDH* (glyceraldehyde-3-phosphate dehydrogenase) gene specific primers (Supplementary Figure S16E, Supplementary Table 1). Further, the fungal colonies were washed with sterile water and sprayed over the rice seedlings after adding two drops of Tween20 surfactant. The development of disease symptoms was further noted (Supplementary Figure S16F).

### Statistical analyses

All the experiments were replicated thrice or more as indicated in the respective figure legends. Standard error of variations (SEVs) of various measurements were represented as error bars. Mean values were compared between the treatments at 0.05 (*) or 0.01 (**) probability levels using Student’s t-test. Non-significant changes were denoted as ‘ns’.

## Acknowledgements

M.M. gratefully acknowledges the financial support from SERB, DST, Government of India under National Post-Doctoral Fellowship (NPDF) scheme (File number: PDF/2020/000511) and project grant from Department of Biotechnology (DBT), Government of India under the BioCARe Women Scientist scheme (File number: BT/PR51354/BIC/101/1344/2023). S.J. acknowledges DBT, Government of India for fellowship. U.P. and G.B. acknowledges Council of Scientific and Industrial Research (CSIR), Government of India for fellowship. A.K.S. acknowledges Sir J.C. Bose National Fellowship Award from SERB, Government of India. Authors also thank the Confocal Microscopy Facility and the Central Instrumental Facility of NIPGR, New Delhi, India. The authors are thankful to DBT-eLibrary Consortium (DELCON) for providing access to e-resources.

## Author contributions

M.M. and A.K.S. conceived the project. M.M. designed the experiments, carried out most of the experiments, analyzed all the data and wrote the manuscript. B.R. developed most the transgenic rice lines, performed microscopy experiments and analyzed photosynthetic parameters. S.J. performed the *in-vitro* kinase assay. U.P. conducted the pull-down assay. G.B. developed and screened the *mpk3* knockout lines. A.K.S. supervised the work, proof-read the manuscript and provided financial support.

## Competing interests

The authors declare no competing interests.

## Additional information

### Supplementary information

Provided as a separate file.

## Notes

### Competing Interest Statement

The authors have declared no competing interest.

## References

Baba, A. I., Andrási, N., Valkai, I., Gorcsa, T., Koczka, L., Darula, Z., Medzihradszky, K. F., Szabados, L., Fehér, A., Rigó, G. and Cséplő, Á. (2019b) AtCRK5 protein kinase exhibits a regulatory role in hypocotyl hook development during skotomorphogenesis. Int. J. Mol. Sci. 20.

Baba, A. I., Valkai, I., Labhane, N. M., Koczka, L., Andrási, N., Klement, É., Darula, Z., Medzihradszky, K. F., Szabados, L., Fehér, A., Rigó, G. and Cséplő, Á. (2019a) CRK5 protein kinase contributes to the progression of embryogenesis of *Arabidopsis thaliana*. Int. J. Mol. Sci. 20.

Barbosa, I. C. and Schwechheimer, C. (2014) Dynamic control of auxin transport-dependent growth by AGCVIII protein kinases. Curr. Opin. Plant Biol. 22.

Bisen, K., Biswas, S. K., Kumar, V., Lal, K., Kumar, R. and Kumar, N. (2015) Biochemical changes in relation to brown leaf spot (*Drechslera oryzae*) resistance in different rice genotypes. J. Plant Stud. 4.

Bradford, M. M. (1976) A rapid and sensitive method for the quantitation of microgram quantities of protein utilizing the principle of protein-dye binding. Anal Biochem. 72.

Canher, B., Heyman, J., Savina, M., Devendran, A., Eekhout, T., Vercauteren, I., Prinsen, E., Matosevich, R., Xu, J., Mironova, V. and De Veylder, L. (2020) Rocks in the auxin stream: Wound-induced auxin accumulation and *ERF115* expression synergistically drive stem cell regeneration. Proc. Natl. Acad. Sci. USA 117.

Chu, J., Xu, H., Dong, H. and Xuan, Y. H. (2021) Loose Plant Architecture 1-Interacting Kinesin-like Protein KLP Promotes Rice Resistance to Sheath Blight Disease. Rice 14.

Dhonukshe, P., Huang, F., Galvan-Ampudia, C. S., Mähönen, A. P., Kleine-Vehn, J., Xu, J., Quint, A., Prasad, K., Friml, J., Scheres, B. and Offringa, R. (2010) Plasma membrane-bound AGC3 kinases phosphorylate PIN auxin carriers at TPRXS(N/S) motifs to direct apical PIN recycling. Development 137.

Ditengou, F. A., Teale, W. D., Kochersperger, P., Flittner, K. A., Kneuper, I., van der Graaff, E., Nziengui, H., Pinosa, F., Li, X., Nitschke, R., Laux, T. and Palme, K. (2008) Mechanical induction of lateral root initiation in *Arabidopsis thaliana*. Proc. Natl. Acad. Sci. USA 105.

Dory, M., Hatzimasoura, E., Kállai, B. M., Nagy, S. K., Jäger, K., Darula, Z., Nádai, T. V., Mészáros, T., López-Juez, E., Barnabás, B., Palme, K., Bögre, L., Ditengou, F. A. and Dóczi, R. (2018) Coevolving MAPK and PID phosphosites indicate an ancient environmental control of PIN auxin transporters in land plants. FEBS Lett. 592.

Fujihira, K., Kurata, T., Watahiki, M. K., Karahara, I. and Yamamoto, K. T. (2000) An agravitropic mutant of Arabidopsis, endodermalamyloplast less1, that lacks amyloplasts in hypocotyl endodermal cell layer. Plant Cell Physiol. 41.

Fukaki, H., Wysocka-Diller, J., Kato, T., Fujisawa, H., Benfey, P.N. and Tasaka, M. (1998) Genetic evidence that the endodermis is essential for shoot gravitropism in *Arabidopsis thaliana*. Plant J. 14.

Grones, P., Abas, M., Hajný, J., Jones, A., Waidmann, S., Kleine-Vehn, J. and Friml, J. (2018) PID/WAG-mediated phosphorylation of the Arabidopsis PIN3 auxin transporter mediates polarity switches during gravitropism. Sci Rep. 8.

Hajný, J., Prát, T., Rydza, N., Rodriguez, L., Tan, S., Verstraeten, I., Domjan, D., Mazur, E., Smakowska-Luzan, E., Smet, W., Mor, E., Nolf, J., Yang, B., Grunewald, W., Molnár, G., Belkhadir, Y., De Rybel, B. and Friml, J. (2020) Receptor kinase module targets PIN-dependent auxin transport during canalization. Science 370.

Harmoko, R., Yoo, J. Y., Ko, K. S., Ramasamy, N. K., Hwang, B. Y., Lee, E. J., Kim, H. S., Lee, K. J., Oh, D. B., Kim, D. Y., Lee, S., Li, Y., Lee, S. Y. and Lee, K. O. (2016) N-glycan containing a core α1,3-fucose residue is required for basipetal auxin transport and gravitropic response in rice (*Oryza sativa*). New Phytol. 212.

Hilgarth, R. S. and Lanigan, T. M. (2019) Optimization of overlap extension PCR for efficient transgene construction. MethodsX 7.

Hoermayer, L., Montesinos, J. C., Marhava, P., Benková, E., Yoshida, S. and Friml, J. (2020) Wounding-induced changes in cellular pressure and localized auxin signalling spatially coordinate restorative divisions in roots. Proc. Natl. Acad. Sci. USA 117.

Hu, Y., Li, S., Fan, X., Song, S., Zhou, X., Weng, X., Xiao, J., Li, X., Xiong, L., You, A. and Xing, Y. (2020) *OsHOX1* and *OsHOX28* Redundantly Shape Rice Tiller Angle by Reducing *HSFA2D* Expression and Auxin Content. Plant Physiol. 184.

Huang, L., Wang, W., Zhang, N., Cai, Y., Liang, Y., Meng, X., Yuan, Y., Li, J., Wu, D., and Wang, Y. (2021). LAZY2 controls rice tiller angle through regulating starch biosynthesis in gravity-sensing cells. New Phytol. 231.

Jatoi, G. H., Keerio, A. U., Abdulle, Y. A. and Qiu, D. (2019) Effect of selected fungicides and Bio-Pesticides on the mycelial colony growth of the *Helminthosporium oryzae*. brown spot of rice. Acta Ecol. Sin. 39.

Jia, W., Li, B., Li, S., Liang, Y., Wu, X., Ma, M., Wang, J., Gao, J., Cai, Y., Zhang, Y., Wang, Y., Li, J. and Wang, Y. (2016) Mitogen-Activated Protein Kinase Cascade MKK7-MPK6 Plays Important Roles in Plant Development and Regulates Shoot Branching by Phosphorylating PIN1 in Arabidopsis. PLoS Biol. 14.

Jonwal, S., Rengasamy, B. and Sinha, A.K. (2023) Regulation of photosynthesis by mitogen-activated protein kinase in rice: antagonistic adjustment by OsMPK3 and OsMPK6. Physiol. Mol. Biol. Plants 29.

Karami, O., Philipsen, C., Rahimi, A., Nurillah, A. R., Boutilier, K. and Offringa, R. (2023) Endogenous auxin maintains embryonic cell identity and promotes somatic embryo development in Arabidopsis. Plant J. 113.

Kircher, S. and Schopfer, P. (2016) Priming and positioning of lateral roots in Arabidopsis. An approach for an integrating concept. J. Exp. Bot. 67.

Lanassa Bassukas, A. E., Xiao, Y. and Schwechheimer, C. (2022) Phosphorylation control of PIN auxin transporters. Curr. Opin. Plant Biol. 65.

Li, H., Sun, H., Jiang, J., Sun, X., Tan, L., and Sun, C. (2021). TAC4 controls tiller angle by regulating the endogenous auxin content and distribution in rice. Plant Biotechnol. J. 19.

Lim, P. O. and Nam, H. G. (2005) The molecular and genetic control of leaf senescence and longevity in Arabidopsis. Curr. Top. Dev. Biol. 67.

Luschnig, C. and Friml, J. (2024) Over 25 years of decrypting PIN-mediated plant development. Nat. Commun. 15.

Manna, M., Rengasamy, B. and Sinha, A. K. (2024a) Nutrient and Water Availability Influence Rice Physiology, Root Architecture and Ionomic Balance via Auxin Signalling. Plant Cell Environ. doi: 10.1111/pce.15171.

Manna, M., Rengasamy, B. and Sinha, A. K. (2024c) A rapid and robust colorimetric method for measuring relative abundance of auxins in plant tissues. Phytochem Anal. 35.

Manna, M., Rengasamy, B., Ambasht, N. K., Sinha and A. K. (2022) Characterization and expression profiling of PIN auxin efflux transporters reveal their role in developmental and abiotic stress conditions in rice. Front Plant Sci. 13.

Manna, M., Rengasamy, B., Reddy, M. K. and Sinha, A. K. (2024b) Revisiting rice transformation for a fail-safe protocol and its application for various gene functional and molecular studies. J. Plant Growth Regul. doi: 10.1007/s00344-024-11486-6.

Mehra, P., Pandey, B. K., Melebari, D., Banda, J., Leftley, N., Couvreur, V., Rowe, J., Anfang, M., De Gernier, H., Morris, E., Sturrock, C. J., Mooney, S. J., Swarup, R., Faulkner, C., Beeckman, T., Bhalerao, R. P., Shani, E., Jones, A. M., Dodd, I. C., Sharp, R. E., Sadanandom, A., Draye, X. and Bennett, M. J. (2022) Hydraulic flux-responsive hormone redistribution determines root branching. Science 378.

Melnyk, C. W. (2016) Plant grafting: insights into tissue regeneration. Regeneration 4.

Morita, M.T. and Tasaka, M. (2024) Gravity sensing and signaling. Curr. Opin. Plant Biol. 7.

Pellegrini-Calace, M., Maiwald, T. and Thornton, J. M. (2009) PoreWalker: a novel tool for the identification and characterization of channels in transmembrane proteins from their three-dimensional structure. PLoS Comput Biol. 5.

Qi, T., Yang, W., Hassan, M. J., Liu, J., Yang, Y., Zhou, Q., Li, H. and Peng, Y. (2024) Genome-wide identification of Aux/IAA gene family in white clover (*Trifolium repens* L.) and functional verification of *TrIAA18* under different abiotic stress. BMC Plant Biol. 24.

Rigó, G., Ayaydin, F., Tietz, O., Zsigmond, L., Kovács, H., Páy, A., Salchert, K., Darula, Z., Medzihradszky, K. F., Szabados, L., Palme, K., Koncz, C. and Cséplo, A. (2013) Inactivation of plasma membrane-localized CDPK-RELATED KINASE5 decelerates PIN2 exocytosis and root gravitropic response in Arabidopsis. Plant Cell 25.

Sack, F. (1997) Plastids and gravitropic sensing. Planta 203.

Savatin, D. V., Gramegna, G., Modesti, V. and Cervone, F. (2014) Wounding in the plant tissue: the defense of a dangerous passage. Front. Plant Sci. 5.

Singh, P., Ara, H., Tayyeba, S., Pandey, C. and Sinha, A. K. (2019) Development of efficient protocol for rice transformation overexpressing MAP kinase and their effect on root phenotypic traits. Protoplasma 256(4).

Su, Y. H., Liu, Y.B. and Zhang, X. S. (2011). Auxin–Cytokinin Interaction Regulates Meristem Development. Mol. Plant. 4.

Sun, Q., Li, T. Y., Li, D. D., Wang, Z. Y., Li, S., Li, D. P., Han, X., Liu, J. M. and Xuan, Y. H. (2019) Overexpression of Loose Plant Architecture 1 increases planting density and resistance to sheath blight disease via activation of PIN-FORMED 1a in rice. Plant Biotechnol J. 17.

Suzuki, H., Kato, H., Iwano, M., Nishihama, R. and Kohchi, T. (2023) Auxin signaling is essential for organogenesis but not for cell survival in the liverwort *Marchantia polymorpha*. Plant Cell 35.

Tivendale, N. D. and Millar, A. H. (2022) How is auxin linked with cellular energy pathways to promote growth? New Phytol. 233.

Ung, K. L., Winkler, M., Schulz, L., Kolb, M., Janacek, D. P., Dedic, E., Stokes, D. L., Hammes, U. Z. and Pedersen, B. P. (2022) Structures and mechanism of the plant PIN-FORMED auxin transporter. Nature 609.

Vega-Muñoz, I., Duran-Flores, D., Fernández-Fernández, Á. D., Heyman, J., Ritter, A. and Stael, S. (2020) Breaking Bad News: Dynamic Molecular Mechanisms of Wound Response in Plants. Front. Plant Sci. 11.

Vidhyasekaran, P., Borromeo, E. S. and Mew, T. W. (1992) *Helminthosporium oryzae* toxin suppresses phenol metabolism in rice plants and aids pathogen colonization. Physiol. Mol. Plant Pathol. 41.

Wang, W., Gao, H., Liang, Y., Li, J. and Wang, Y. (2022) Molecular basis underlying rice tiller angle: Current progress and future perspectives. Mol. Plant 15.

Xia, X., Mi, X., Jin, L., Guo, R., Zhu, J., Xie, H., Liu, L., An, Y., Zhang, C., Wei, C. and Liu, S. (2021) CsLAZY1 mediates shoot gravitropism and branch angle in tea plants (*Camellia sinensis*). BMC Plant Biol. 21.

Xu, L. (2018) De novo root regeneration from leaf explants: wounding, auxin, and cell fate transition. Curr. Opin. Plant Biol. 41.

Xu, M., Zhu, L., Shou, H. and Wu, P. A. (2005) PIN1 family gene, OsPIN1, involved in auxin-dependent adventitious root emergence and tillering in rice. Plant Cell Physiol. 46.

Yu, Z., Zhang, F., Friml, J. and Ding, Z. (2022) Auxin signaling: Research advances over the past 30 years. J. Integr. Plant Biol. 64(2).

Zhang, J., Nodzynski, T., Pencík, A., Rolcík, J. and Friml, J. (2010) PIN phosphorylation is sufficient to mediate PIN polarity and direct auxin transport. Proc. Natl. Acad. Sci. USA 107.

Zhang, N., Yu, H., Yu, H., Cai, Y., Huang, L., Xu, C., Xiong, G., Meng, X., Wang, J., Chen, H., Liu, G., Jing, Y., Yuan, Y., Liang, Y., Li, S., Smith, S. M., Li, J. and Wang, Y. A. (2018) Core Regulatory Pathway Controlling Rice Tiller Angle Mediated by the *LAZY1*-Dependent Asymmetric Distribution of Auxin. Plant Cell 30.

Zourelidou, M., Absmanner, B., Weller, B., Barbosa, I. C., Willige, B. C., Fastner, A., Streit, V., Port, S. A., Colcombet, J., de la Fuente van Bentem, S., Hirt, H., Kuster, B., Schulze, W. X., Hammes, U. Z. and Schwechheimer, C. (2014) Auxin efflux by PIN-FORMED proteins is activated by two different protein kinases, D6 PROTEIN KINASE and PINOID. Elife 3.

Zwiewka, M., Bilanovičová, V., Seifu, Y. W. and Nodzyński T. (2019) The Nuts and Bolts of PIN Auxin Efflux Carriers. Front. Plant Sci. 10.

