## Supplementary Material for "A positive feedback loop of ‘MPK3-PIN1A trafficking-auxin flux’ trio governs dual gravitropic and wounding response in rice"

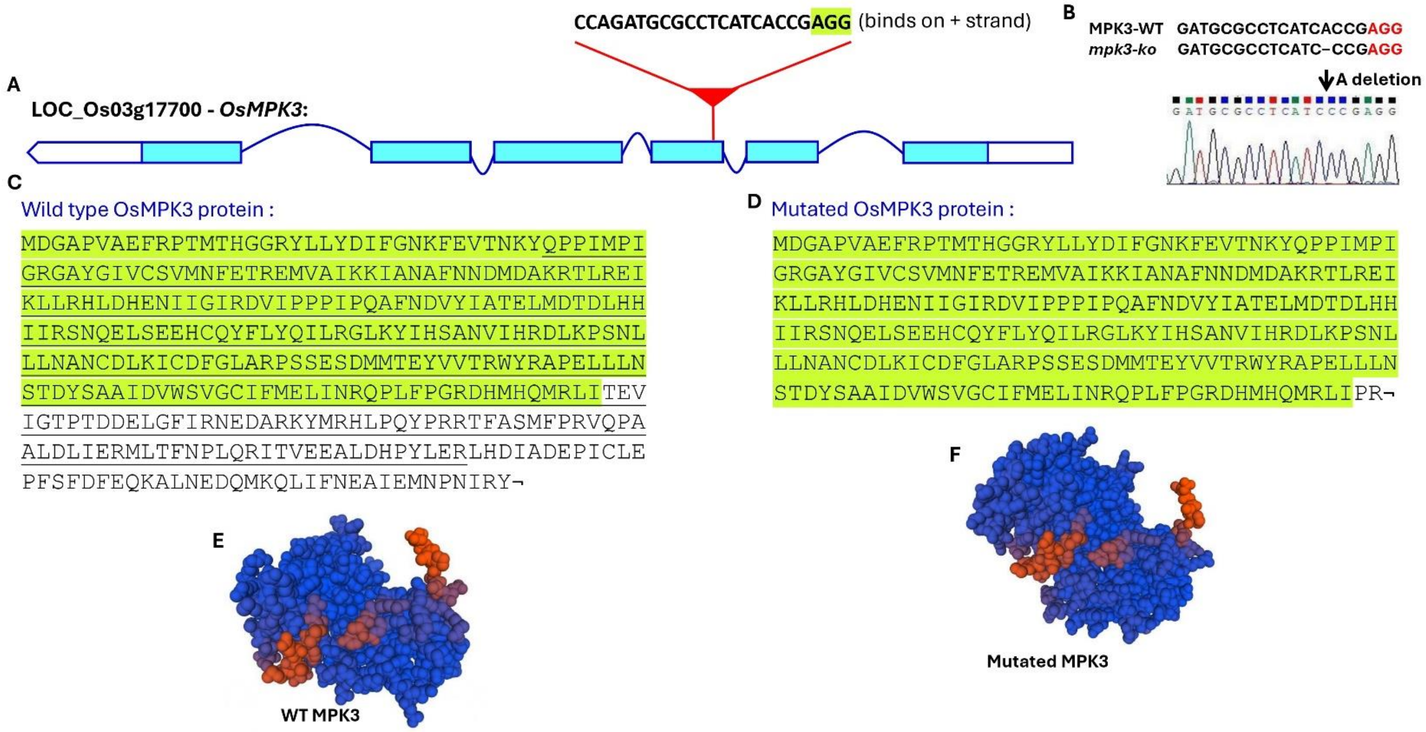


**Supplementary Figure S1. sgRNA-Cas9 mediated knockout of *mpk3* gene in rice. (A)** Schematic representation of *OsMPK3* gene and the sgRNA targeting the fourth exon of the gene. **(B)** Sequencing confirmation of an *mpk3* knockout line showing deletion of base 'A' in the sgRNA binding region of the gene. **(C, D)** Comparison of the wild-type (WT) and truncated MPK3 amino acid sequence (green shaded amino acid sequences depict the extent of similarity between them and the underlined amino acid sequence is the active site of MPK3). **(E, F)** Predicted 3D structures of the WT and truncated MPK3 proteins.

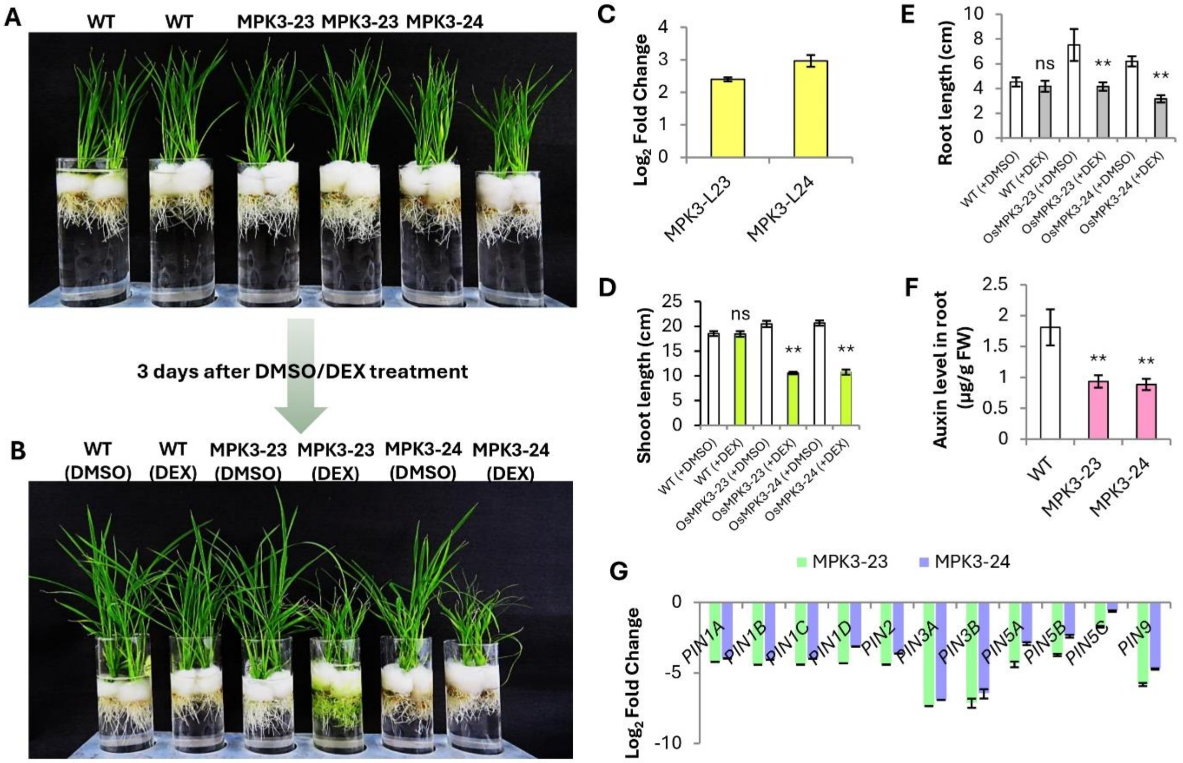


**Supplementary Figure S2. The morphology and physiology of the DEX inducible MPK3 overexpression lines. (A, B)** DEX treatment slowed down the growth rate of *MPK3* overexpression lines; MPK3-23 and MPK3-24. **(C)** Increase in expression of *OsMPK3* gene following DEX treatment as revealed by qRT-PCR (n=3). Increased expression of *MPK3* decreases shoot (n=10) **(D)** and root (n=10) **(E)** growth of the rice seedlings, decrease auxin levels (n=4) **(F)** and downregulate the expression of *PIN* genes (n=3) **(G)**. WT: wild type rice seedlings.

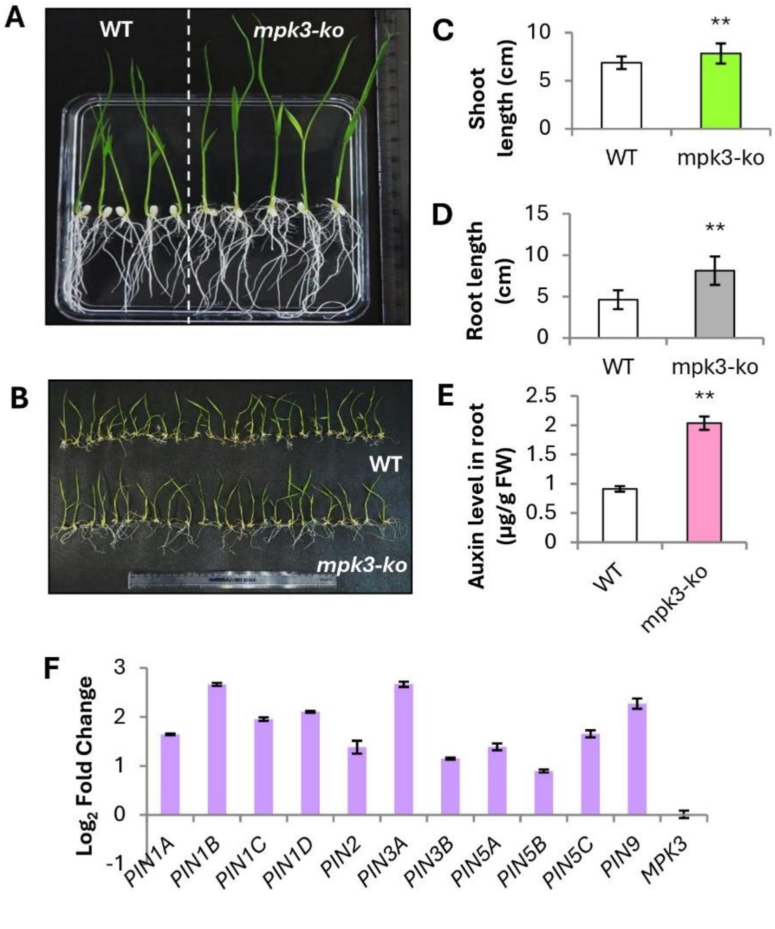


**Supplementary Figure S3. The morphology and physiology of the *mpk3* knockout line.** Comparison of seedling morphology of wild-type (WT) and the mpk3 knockout line (*mpk3-ko*) grown over half-strength solid **(A)** or liquid MS medium **(B)**. The shoot length (n=25) **(C)** and root length (n=25) **(D)** of *mpk3-ko* seedlings were significantly higher than the wild-type (WT) seedlings grown under similar conditions. Moreover, the root auxin content of *mpk3-ko* seedlings was higher than the WT (n=4) **(E)** along with having higher expression of the *PIN* genes (n=3) **(F)**.

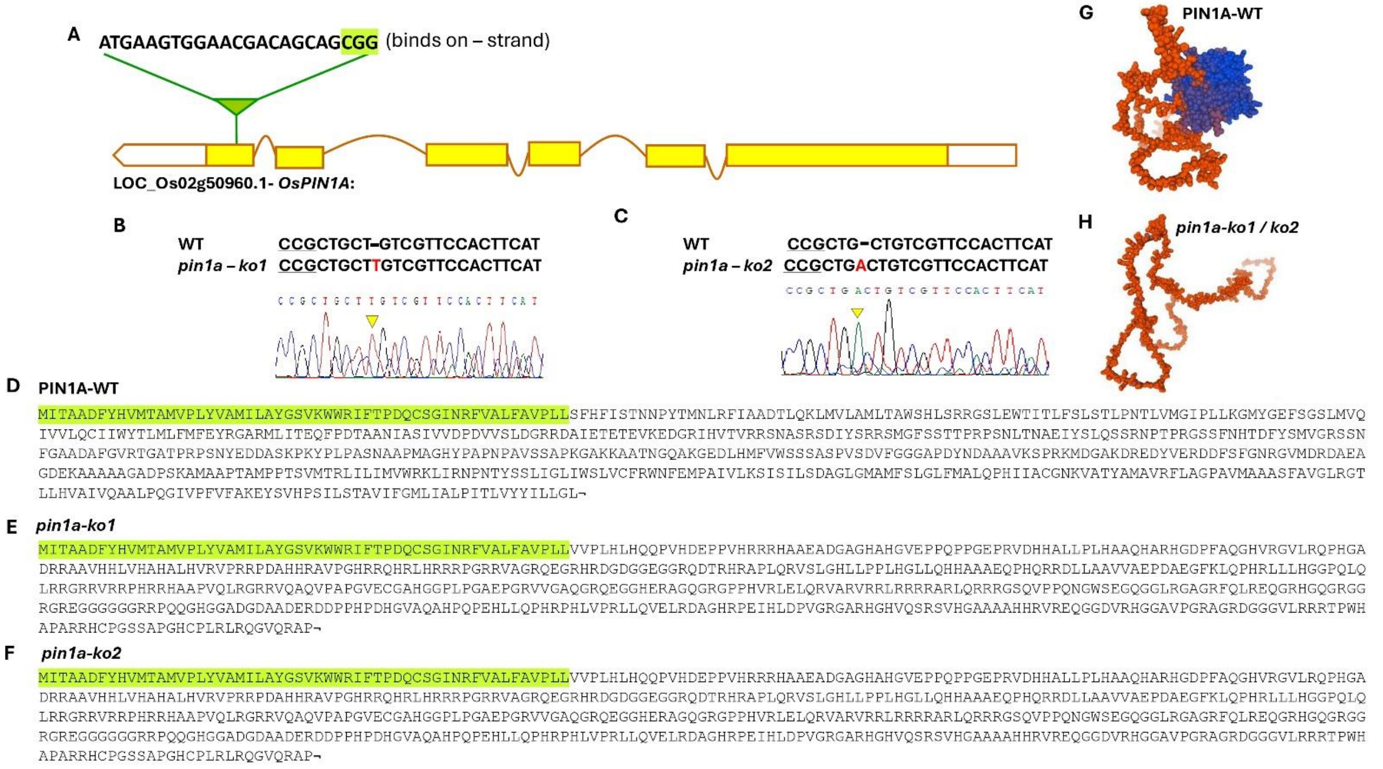


**Supplementary Figure S4. The generation of *pin1a* knockout lines in rice. (A)** Schematic representation of *OsPIN1A* gene and the sgRNA targeting the first exon of the gene. **(B, C)** Sequencing confirmation of the *pin1a* knockout lines (*pin1a-ko1* and *pin1a-ko2*) showing insertion of 'T' or 'A' base in the sgRNA binding region of the gene. **(D-F)** Comparison of the wild-type (WT) and mutated PIN1A amino acid sequence (green shaded amino acid sequences depict the extent of similarity between them). **(G, H)** Predicted 3D structures of the WT and mutated PIN1A proteins.

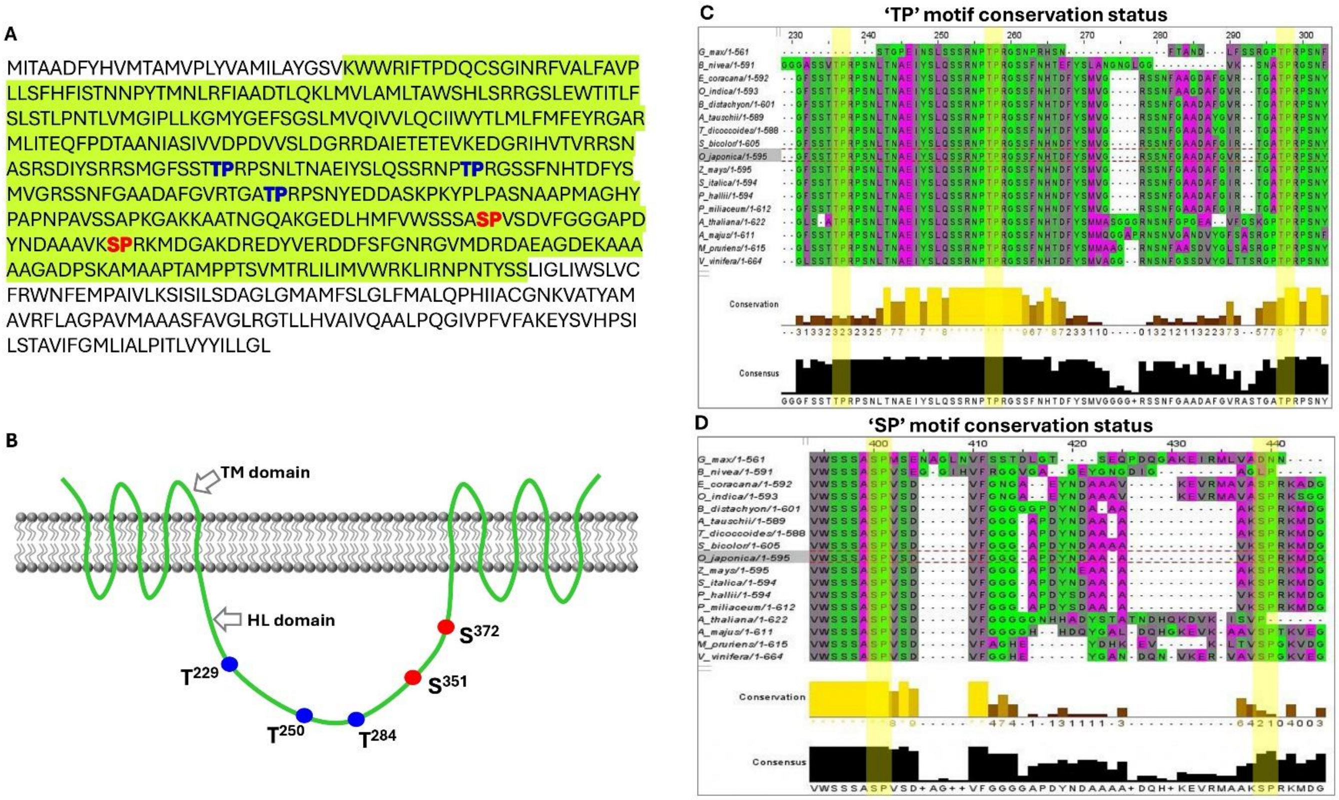


**Supplementary Figure S5. The PIN1A protein sequence and MAPK phosphorylation target sites. (A)** The PIN1A amino acid sequence has five putative MAPK phosphorylation motifs (i.e. three 'TP' and two 'SP' motifs where MAPK phosphorylates the T/S residues). The green shaded portion constitutes the hydrophilic loop (HL) domain, and the flanking portions constitute the trans-membrane (TM) domain. **(B)** The schematic representation of PIN1A protein integrated in the cell membrane. **(C)** The multiple sequence alignment showing the extent of 'TP' motif conservation in PIN1A across the plant species. **(D)** The multiple sequence alignment showing the extent of 'SP' motif conservation in PIN1A across the plant species.

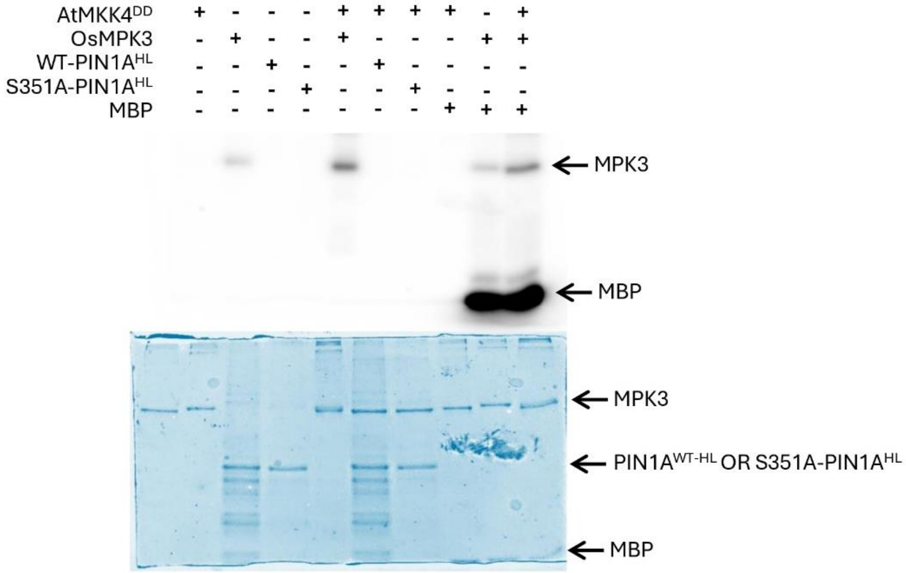


**Supplementary Figure S6. The *in-gel* kinase assay showing activation status of MPK3 protein used in the assay.** The MPK3 protein is auto activated even in the absence of Arabidopsis MKK4$^{DD}$ (lane 2) while in presence of the upstream MAPKK (here Arabidopsis MKK4$^{DD}$), its activity gets enhanced (lane 5). Further, the activated MPK3 was highly functional as it phosphorylated myelin basic protein (MBP) efficiently (lane 9, 10).

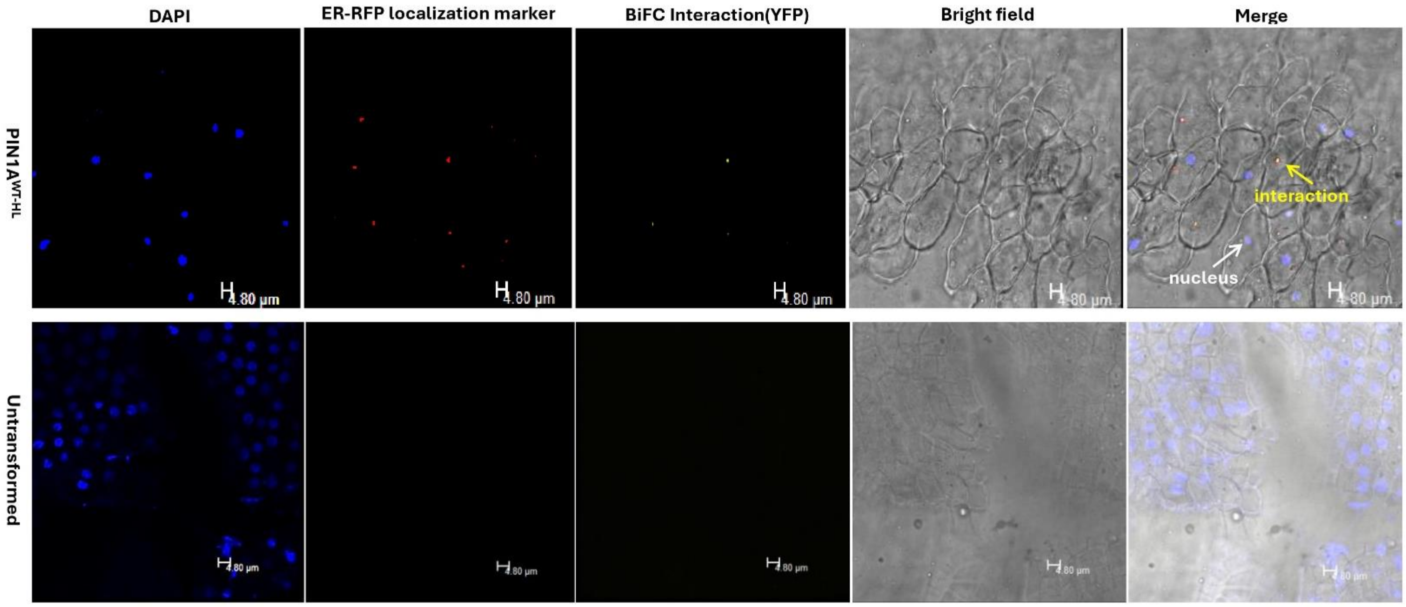


**Supplementary Figure S7. Proving that the PIN1A ferrying transport vesicles are ER remnants or endosomes.** Co-transforming the rice callus cells with BiFC constructs of MPK3 and PIN1A$^{HL}$ (which upon interaction produces YFP signal) and ER-RFP localization marker and subsequently staining the cells with DAPI (highlighted the nuclei of the cells) revealed that YFP signals co localized with the RFP signals thus proving endosomal localization of PIN1A.

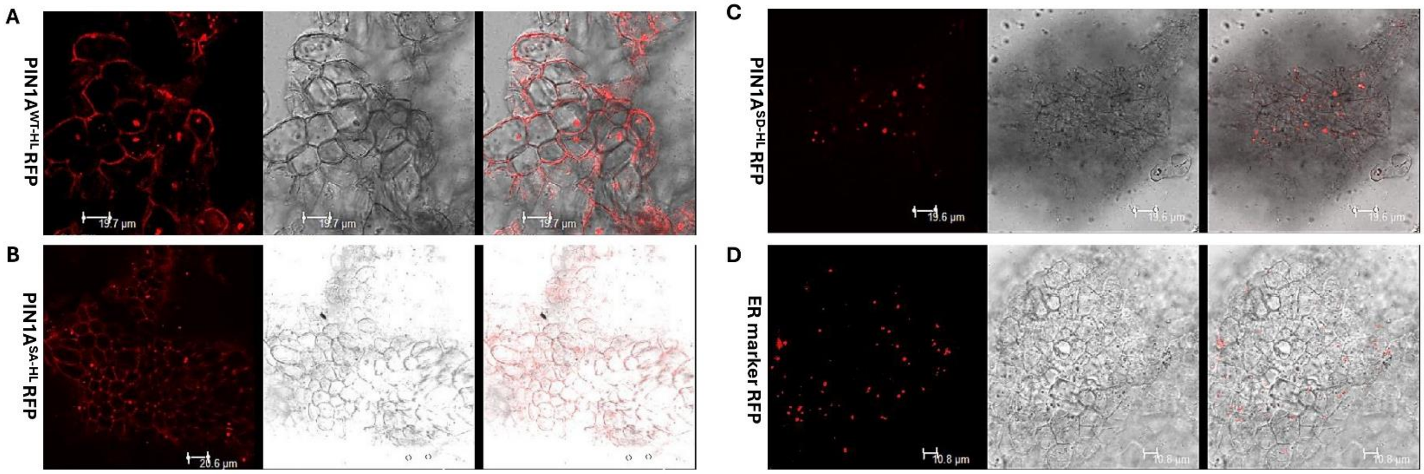


**Supplementary Figure S8. Endosome mediated PIN1A trafficking in rice callus cells.** Transient transformation of rice callus cells with PIN1A$^{WT-HL}$ RFP **(A)**, PIN1A$^{SA-HL}$ RFP **(B)**, PIN1A$^{SD-HL}$ RFP **(C)** and ER-RFP marker **(D)** revealed that all of them had the same pattern of localization signals.

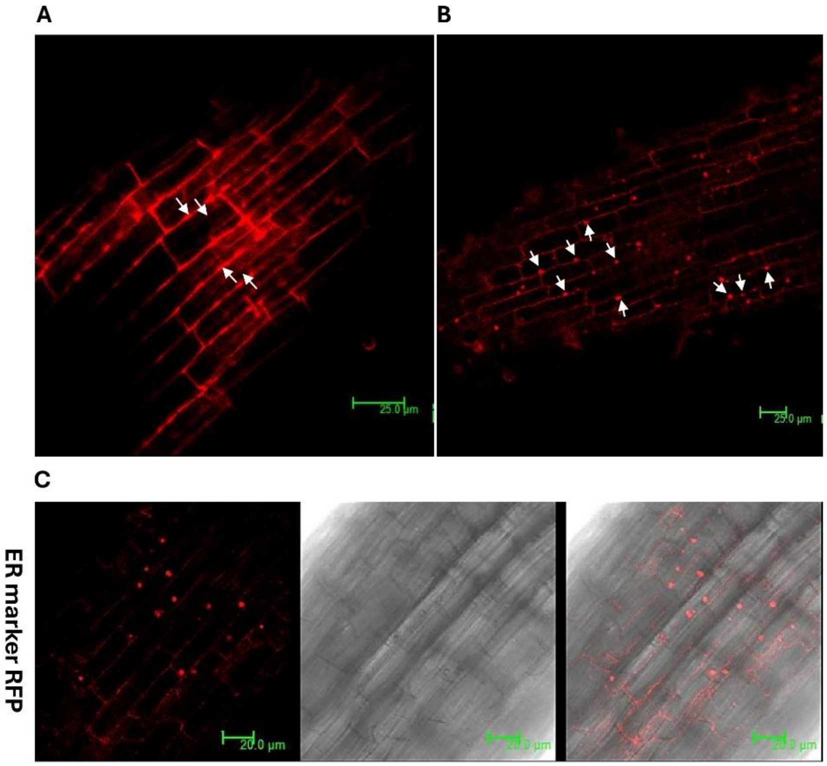


**Supplementary Figure S9. The snapshot view of endosomal vesicle formation from plasma membrane.** Root tissue of the stable transgenic line of PIN1A$^{SD-HL}$ RFP showing protuberances in the plasma membrane **(A)** and subsequent separation of endosomal transport vesicles containing the PIN protein **(B)**. **(C)** Transformation of rice root tissue with ER-RFP localization marker also produces a similar pattern of signal.

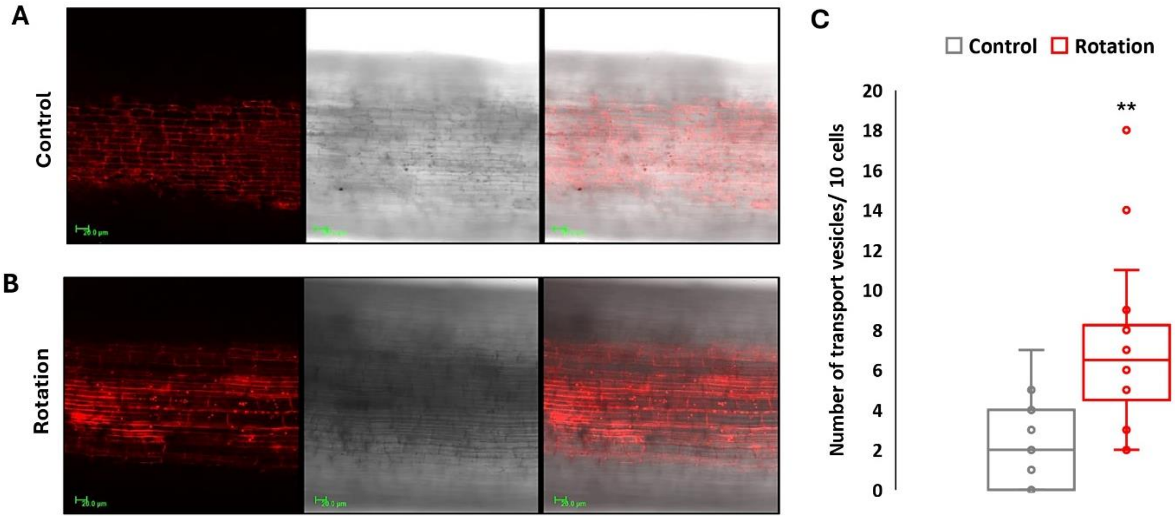


**Supplementary Figure S10. Physical movement in plant organ induces endosomal PIN1A trafficking. (A, B)** Dissected living roots were placed in water in an eppendorf tube and rotated for four hours in a rotor following which they were visualized under the confocal microscope. Dissected living roots kept still in water served as the control. The roots subjected to rotation motion produced more endosomal transport vesicles as compared to the control (n = 10) **(C)**.

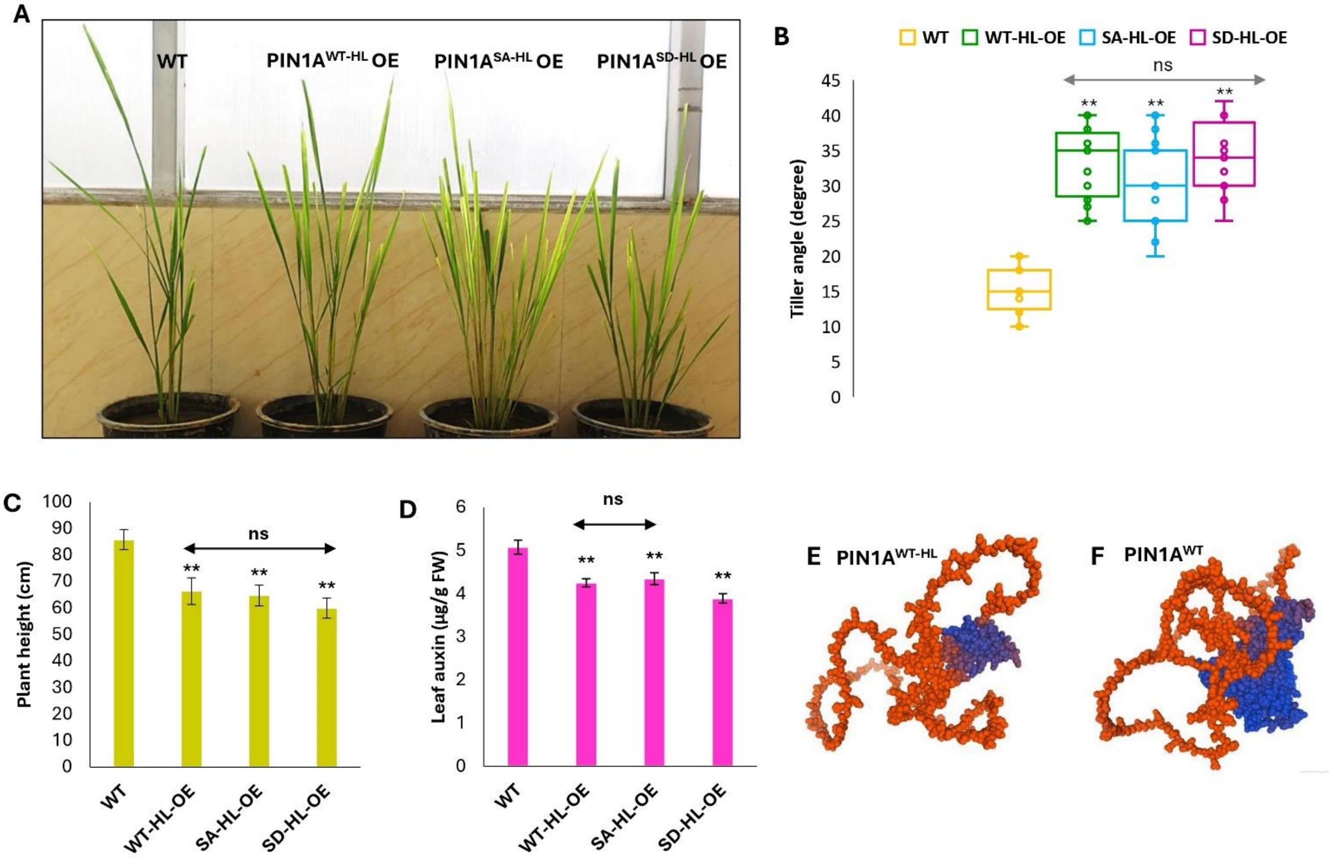


**Supplementary Figure S11. The morphology and physiology of the transgenic rice plants overexpressing HL domain of wild-type/phospho-dead/phospho-mimic PIN1A.** The overexpression lines had more tiller angle (n = 8) **(A, B)**, lesser height (n = 8) **(C)** and less leaf auxin content (n = 4) **(D)**. **(E, F)** The predicted protein structures of only HL domain of PIN1A (PIN1A$^{WT\text{-}HL}$) and full length PIN1A (PIN1A$^{WT}$).

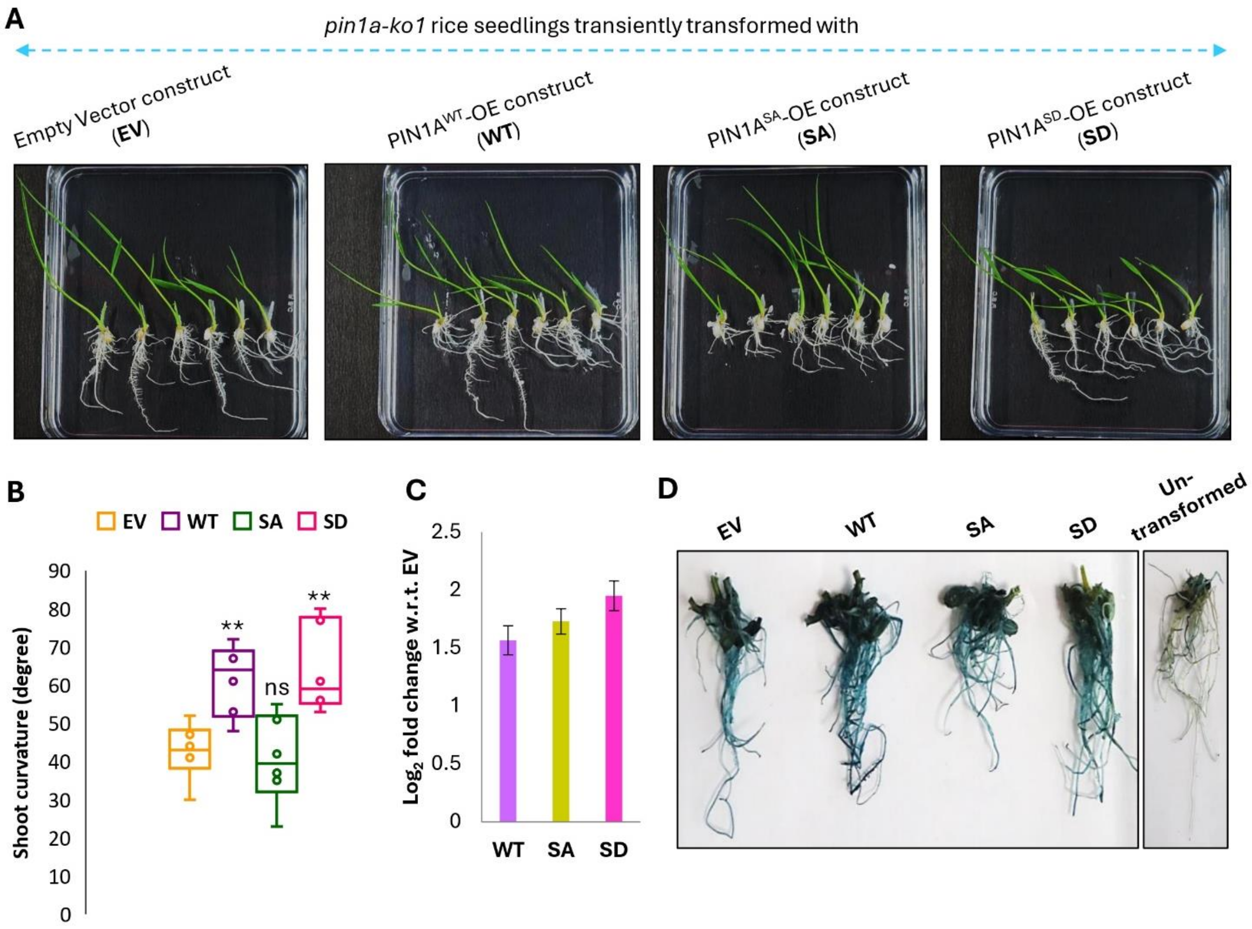


**Supplementary Figure S12. Wild-type and phospho-mimic PIN1A constructs recovered gravitropic response of *pin1a* knockout seedlings. (A)** Analysis of gravitropic response of *pin1a* knockout seedlings after transiently transforming with empty vector (EV)/wild-type (WT)/phospho-dead (SA)/ phospho-mimic (SD) *PIN1A* constructs. For this experiment, the *pin1a* knockout seedlings were incubated in *Agrobacterium* suspensions harboring respective empty vector or *PIN1A* constructs for overnight, following which they were placed straight over half-strength MS medium. The plates were then inverted 90° and shoot curvature was noted after 3 days. **(B)** The wild-type and phospho-mimic *PIN1A* constructs recovered gravitropic response of *pin1a* knockout seedlings while others could not (n = 6). The transformation efficiency was checked by analyzing *PIN1A* expression (n = 3) **(C)** and gus staining (PANIC6b vector contains the reporter marker gene *gus* under control of the Switchgrass ubiquitin promoter) of root tissues **(D)**.

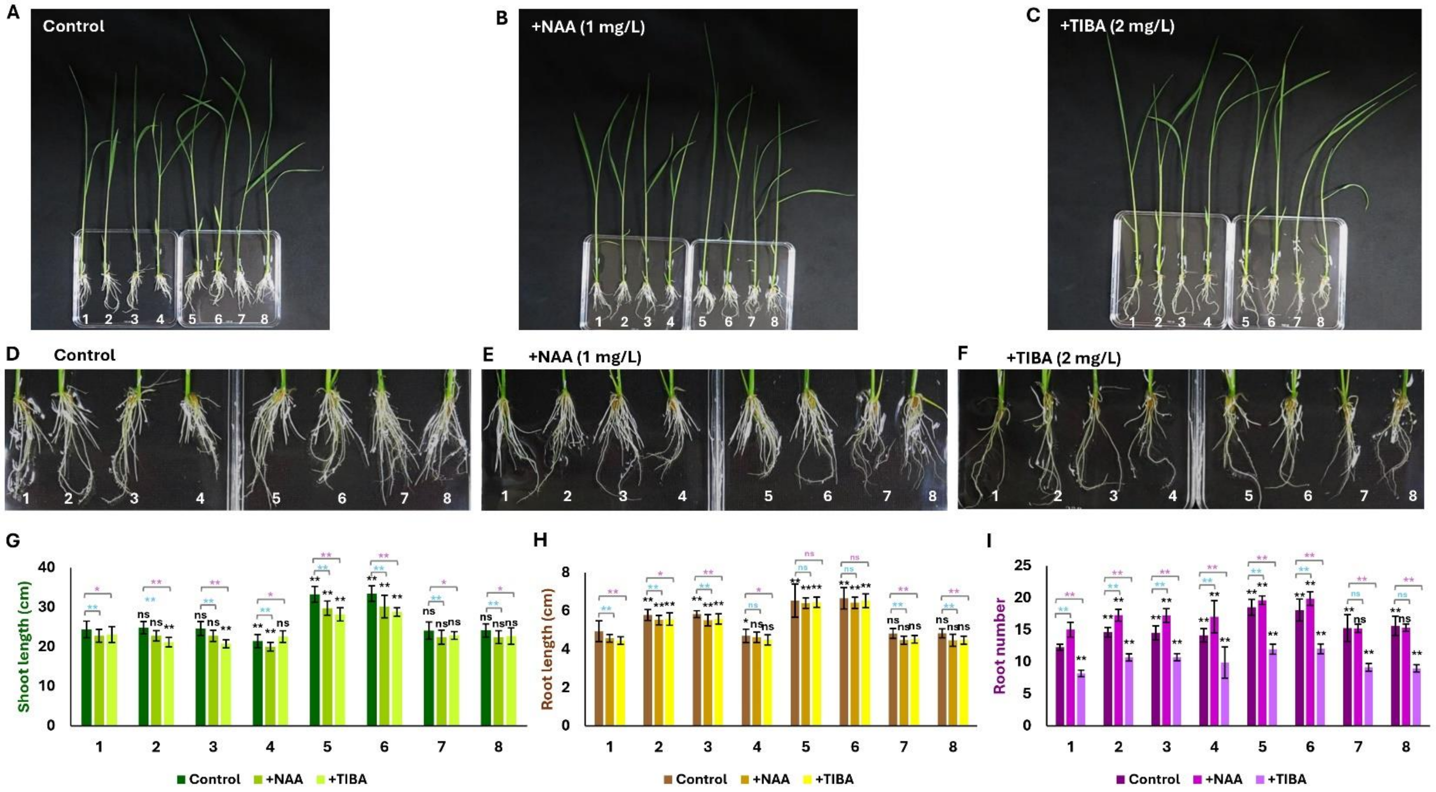


**Supplementary Figure S13. Analysis of seedling morphology of various *PIN1A* overexpression and knockout lines. (A-F)** Whole plant and root morphologies of wild-type [WT; 1], wild-type *PIN1A* overexpression line 1 [2], wild-type *PIN1A* overexpression line 2 [3], phospho-dead *PIN1A* overexpression line [4], phospho-mimic *PIN1A* overexpression line 1 [5], phospho-mimic *PIN1A* overexpression line 2 [6], *pin1a* knockout line 1 [7] and *pin1a* knockout line 2 [8] after growing in one-fifth strength MS medium for two weeks without (control condition) or with NAA/ TIBA. **(G)** Under control condition shoot length of phospho-mimic *PIN1A* overexpression lines were more than WT and shoot length of phospho-dead *PIN1A* overexpression line was shorter than WT. Same was the case in presence of NAA. However, when grown in presence of TIBA, shoot length of phospho-mimic *PIN1A* overexpression lines were more than WT only. **(H)** Under control condition, root lengths of all types of *PIN1A* overexpression lines were larger than WT. However, upon NAA/TIBA treatments, roots of wild-type *PIN1A* overexpression and phospho-mimic *PIN1A* overexpression lines were longer than WT. **(I)** Under control condition, root numbers were more in all the *PIN1A* overexpression and knockout lines. The NAA treatment though increased root formation in all the *PIN1A* overexpression, the same was not evident in the *pin1a* knockout lines. When grown in presence of TIBA, the root formation was more in all the *PIN1A* overexpression lines while root formation was inhibited in the *pin1a* knockout lines. n = 20.

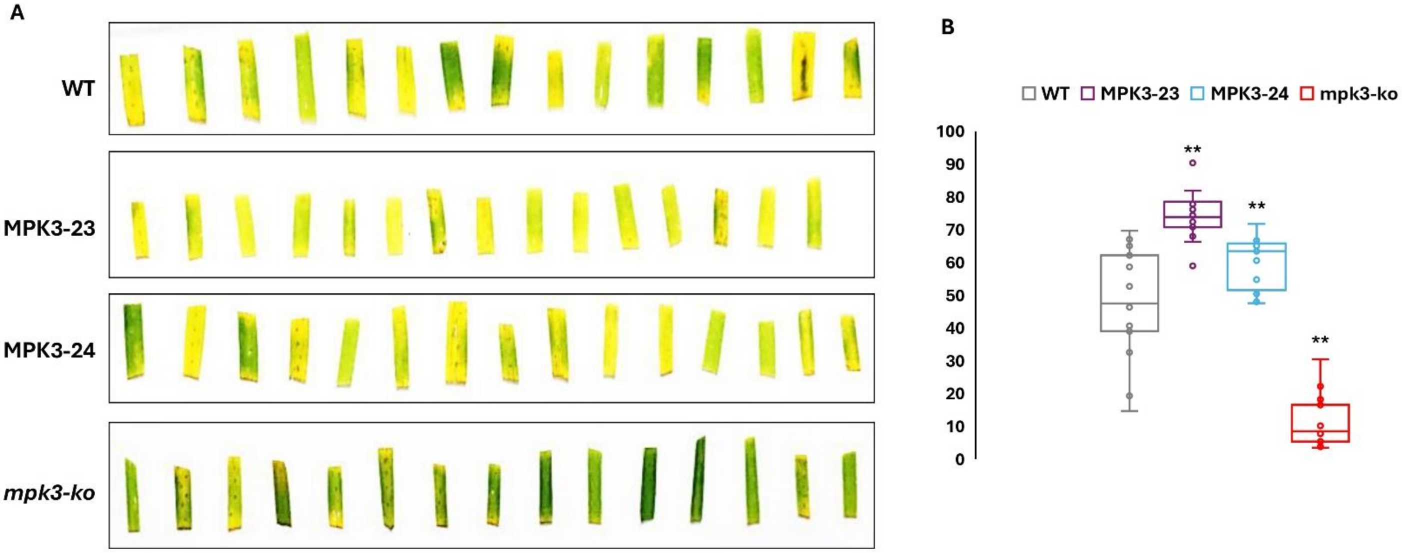


**Supplementary Figure S14. Comparing the wounding response of *MPK3* overexpression and knockout plants.** The phenotype of leaf bits undergoing senescence after wounding **(A)** and quantification of the same revealed that leaves of *mpk3* knockout line (*mpk3-ko*) retained more greenness as compared to wild-type (WT) while the leaves of *MPK3* overexpression lines (MPK3-23 and MPK3-24) turned yellow faster as compared to the WT leaves after wounding (n = 15) **(B)**.

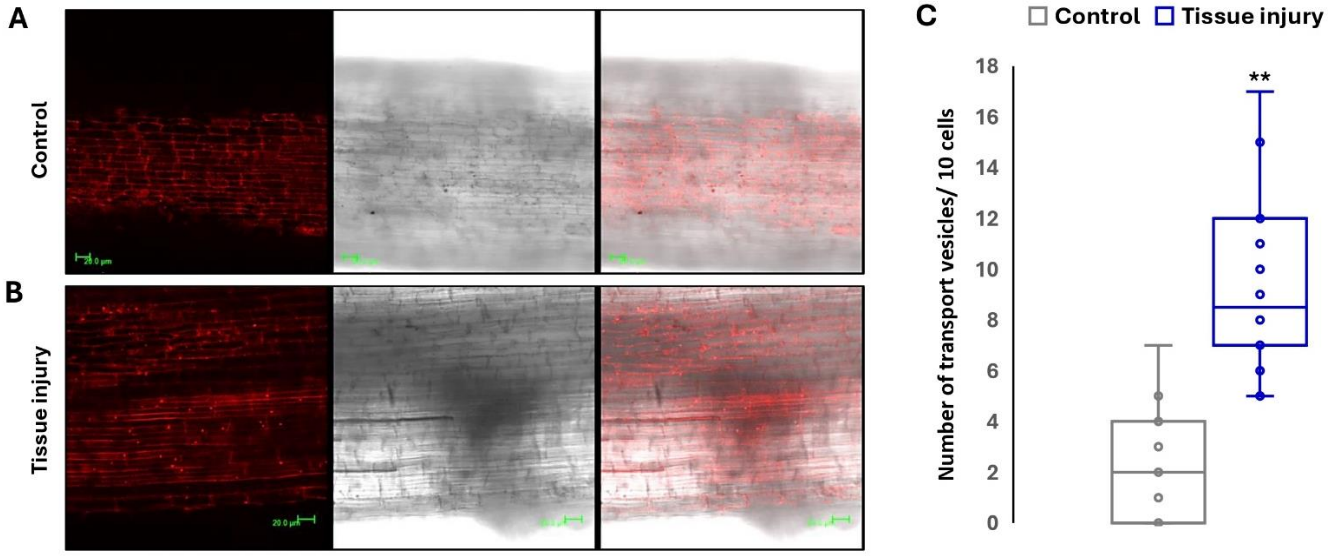


**Supplementary Figure S15. Wounding induces endosomal PIN1A trafficking. (A, B)** Uniformly dissected living roots pieces were fist pricked with a needle (each root bit pricked 10 times) and thereafter placed in water in an eppendorf tube and incubated at room temperature for four hours following which they were visualized under the confocal microscope. Dissected living roots kept still in water served as the control. The wounded roots produced more endosomal transport vesicles as compared to the control (n = 10) **(C)**.

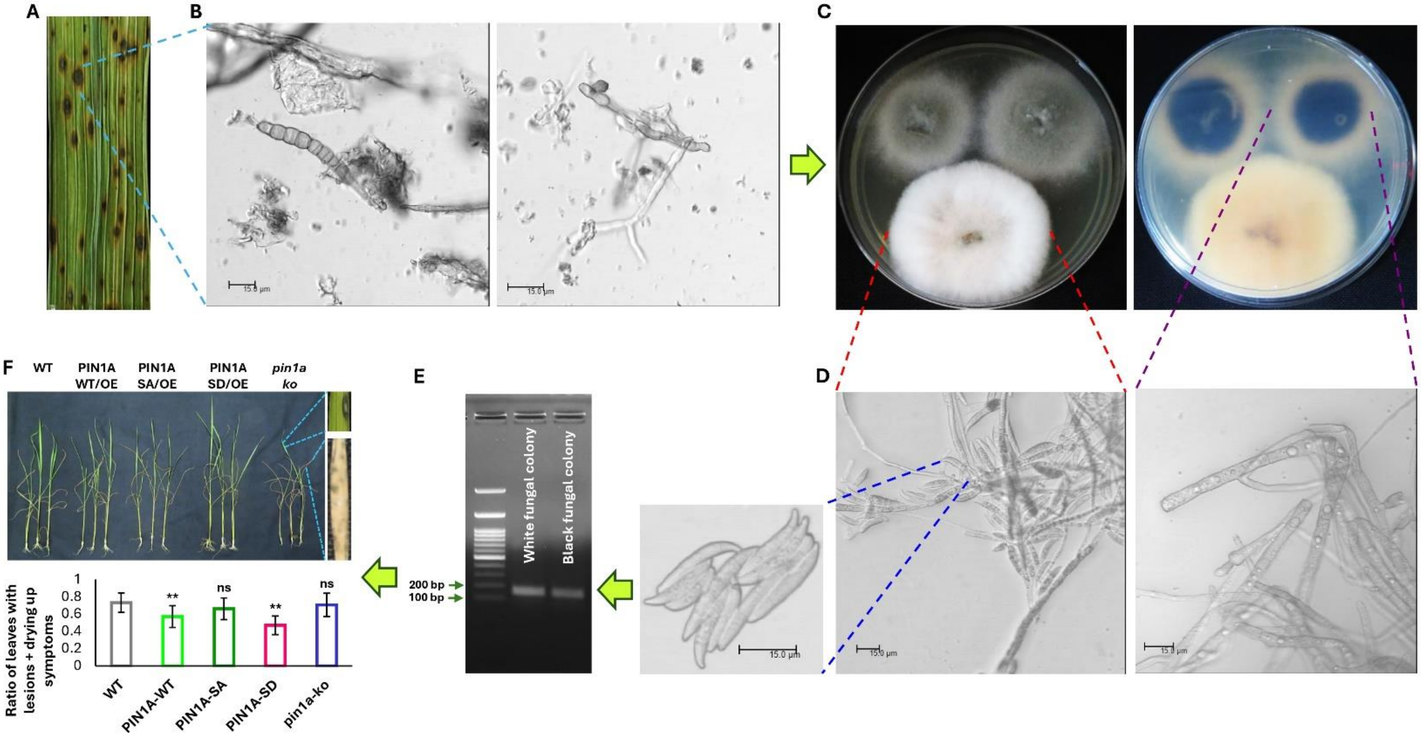


**Supplementary Figure S16. Isolation and characterization of fungal pathogen causing leaf spot disease in rice.** The visualization of leaf spot **(A)** scrapings under the confocal microscope revealed presence of fungal spores of *Bipolaris oryzae* and fungal hyphae **(B)**. To confirm the pathogen, the leaf bits containing diseased lesions were sterilized and incubated over PDA medium which resulted in formation two types of fungal colonies (front and back views are being shown here) **(C)**. **(D)** The white fungal colony was found to produce typical conidia of *Bipolaris oryzae* while sporulation was missing in the black fungal colony. **(E)** Molecular confirmation of genomic DNA of these two types of fungi by PCR with (glyceraldehyde-3-phosphate dehydrogenase) gene specific primers reveled that both the types of fungi were indeed *Bipolaris oryzae*. **(F)** Reinoculation of fungal spores in the wild-type/ *PIN1A* overexpression/ *pin1a*-knockout plants revealed appearance of disease symptoms which was significantly less severe in case of wild-type *PIN1A* overexpression line and phospho-mimic *PIN1A* overexpression line (n = 17).

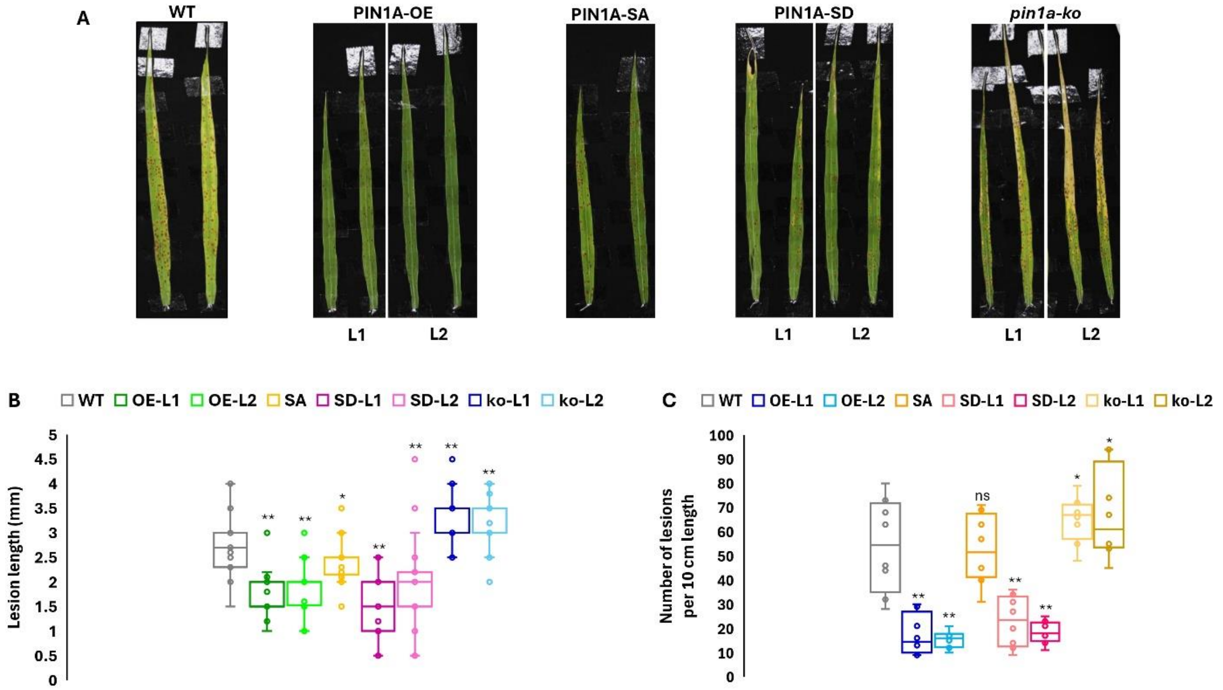


**Supplementary Figure 17.** ***Bipolaris oryzae*** **produces differential brown spot disease symptom in various** ***PIN1A*** **overexpression and knockout lines. (A)** The phenotypic observation of brown spot disease development in flag leaves of respective wild-type (WT)/*PIN1A* overexpression (OE)/ *pin1a* knockout (ko) lines. Quantification of lesion lengths (n = 21-49) **(B)** and lesion density (n = 9) **(C)** revealed that wild-type *PIN1A* overexpression lines (PIN1A-OE) and phospho-mimic *PIN1A* overexpression lines (PIN1A-SD) were more tolerant to the pathogen than WT plants while the disease severity was more in the *pin1a* knockout lines and phospho-dead HL domain overexpressing rice plants.

**Supplementary Table S1:** Primers used in the present study

| Primer name | Forward primer (5' → 3') | Reverse primer (5' → 3') |
|---|---|---|
| *PIN1A_full length_pENTR* | CACCATGATTACGGCGGCG | TTACAGCCCAAGCAAGATGTAG |
| *PIN1A(HL)_pENTR* | CACCTTCGAGTACCGCGGCGC | GCTGGAGTAGGTGTTCGGG |
| *PIN1A(HL)_pET28a* | ATTACATATGTTCGAGTACCGCGGCGC | AATTGAATTCGCTGGAGTAGGTGTTCGGG |
| *PIN1A_internal_S>A*$^{351}$ | TCCAGCGCGGCTCCCGTGT | ACACGGGAGCCGCGCTGGA |
| *PIN1A_internal_S>D*$^{351}$ | TCCAGCGCGGATCCCGTGT | ACACGGGATCCGCGCTGGA |
| *PIN1A_gRNA* | GGCAATGAAGTGGAACGACAGCAG | AAACCTGCTGTCGTTCCACTTCAT |
| *PIN1A_knockout_ screening* | ATGATTACGGCGGCG | TATCCTGCCGTCCTCCTTCA |
| *PIN1A_knockout_ sequencing* | GTGGCGATGATACTGGCGTA | - |
| *MPK3_gRNA* | GGCAGATGCGCCTCATCACCGAGG | AAACCCTCGGTGATGAGGCGCATC |
| *MPK3_knockout_ screening* | CGCTCCAACCAAGAACTGTCAG | GGTGAGCATCCTCTCGATGAG |
| *MPK3_knockout_ sequencing* | CAACTCCACCGACTACTCC | - |
| *TAR1_RT* | GAAGGAAGGGGTGGACGAC | AGTTCATGGCGGCGAGG |
| *TAR2_RT* | GGTGCGATAGGGAGGATGTG | CGAGAGGCGGTTGATGAAGA |
| *YUCCA1_RT* | TTGGGACGCTAGACCACATC | CAAGTCACCGGCATCCTTGA |
| *YUCCA4_RT* | ATGGCGTGGAGTTTGTGGAT | GCCTTGAGAAACCAACAGCG |
| *YUCCA8_RT* | TGGTCTCAAGAGGCCCAAAC | TCCGTGAACAACTCACCGTC |
| *PIN1A_RT* | CGTCTGCTTCAGGTGGAACT | GGTGATAGGCAAGGCGATGA |
| *PIN1B_RT* | TGGTCCCTCGTCTCCTACAG | GCAAACACAAAGGGCACGAT |
| *PIN1C_RT* | CTCATCGGCCTCATCTGGTC | AGCCCCAGCAGGATGTAGTA |
| *PIN1D_RT* | AGGTGGGGAATTGAGATGCC | GCCATGGCATACGAAGCAAG |
| *PIN2_RT* | GCGCAAGCTCATCAGAAACC | GCAAATGTCGCAACGGTCTT |
| *PIN3A_RT* | GGCCATGTTTAGCCTGGGAT | TTACCGCTGTGCTCAGGATG |
| *PIN3B_RT* | CCATTCTCTCCGATGCAGGG | GGCACAATTCCTTGTGGCAG |
| *PIN5A_RT* | ATGTCCAAGTCAGGCACAGG | AGCATGCAGCCCGTATTCTT |
| *PIN5B_RT* | GGGTTTGTTCATGGCGTTGC | GAAGCGCAGCCTGTATGATG |
| *PIN5C_RT* | GGGCTTCATGCCGATGTACT | TAGACAAAGCCCAGAACCGC |

| | | |
|---|---|---|
| *PIN9_RT* | GTCATCTGGATGGCGGTGAA | AATGATGTCACTGCCAGGGG |
| *18S_RT* | TTAGGCCACGGAAGTTTGAG | GTACAAAGGGCAGGGACGTA |
| *MPK3_RT* | GCTCCAACCAAGAACTGTC | AGTCGCAGA TCTTGAGG |
| *Bipolaris oryzae_ GAPDH* | CGAGTCTACCGGTGTCTTCA | ACCTCAATGTCGGGCTTGTA |
